# Population-specific bycatch risks in two vulnerable anadromous clupeids: insights from otolith microchemistry

**DOI:** 10.1101/2025.05.23.655712

**Authors:** David J. Nachón, Alejando Pico-Calvo, Françoise Daverat, Rufino Vieira-Lanero, Rosa M. Crujeiras, Ana F. Belo, Catarina S. Mateus, Bernardo Quintella, Pedro R. Almeida, Carlos Antunes, Gilles Bareille, Cristophe Pécheyran, Fanny Claverie, Patrick Lambert, Géraldine Lasalle, Fernando Cobo

**Affiliations:** Departamento de Zooloxía e Antropoloxía Física, Facultade de Bioloxía, Universidade de Santiago de Compostela, Campus Vida, C/Lope Gómez de Marzoa, s/n, 15782, Santiago de Compostela; UMR ECOBIOP, 1224, Institut National de Recherche pour l’Agriculture, l’alimentation et l’Environnement—INRAE, UPPA, 173 Route de Saint-Jean-de-Luz, Saint-Pée-sur-Nivelle 64310, France; Galician Center for Mathematical Research and Technology, CITMA, Universidade de Santiago de Compostela, Campus Vida, 15782, Santiago de Compostela, Spain; MARE—Marine and Environmental Sciences Centre/ARNET—Aquatic Research Network, Universidade de Evora, Evora, Portugal; Departmento de Biologia, Universidade de Évora, Évora, Portugal; Departamento de Biologia Animal, Facultade Ciências, Universidade de Lisboa/MARE—Marine and Environmental Sciences Centre/ARNET—Aquatic Research Network; CIMAR/CIIMAR—Centro Interdisciplinar de Investigação Marinha e Ambiental, Universidade do Porto, 4450-208 Matosinhos, Portugal; Aquamuseu do Rio Minho, 4920-290 Vila Nova de Cerveira, Portugal; University of Pau and the Adour Region, E2S UPPA, CNRS, IPREM, MIRA, Pau, France; Institut National de Recherche pour l’Agriculture, l’alimentation et l’Environnement— INRAE, UR EABX, 33612 Cestas, France

**Keywords:** *Alosa* spp., bycatch hotspots, Iberian coast, metapopulation dynamics, natal origin, trace markers

## Abstract

Otolith microchemistry analysis revealed that bycatch of European shads—allis shad *Alosa alosa* (L. 1758) and twaite shad *Alosa fallax* (Lacépède 1803)— in western Iberian commercial fisheries removes individuals from a wide array of natal origins, with the most abundant source rivers suffering the heaviest losses. Spatial variation in bycatch risk was evident: specific marine areas exhibited high natal-origin diversity, reflecting complex dispersal. *A. alosa* showed extensive medium- and long-distance movements— including rare longitudinal displacements along the Cantabrian slope—and greater natal origin diversity than *A. fallax*, whose dispersal was largely restricted to middle-distance, latitudinal migrations. In both species, bycatches were dominated by the most abundant continental populations—Mondego and Minho for *A. alosa*, and Ulla and Minho for *A. fallax*—suggesting that these rivers function as source populations exporting individuals to sink populations through marine dispersal. Despite their differing dispersal ranges, both species displayed dual resident–dispersive contingents coexisting within the same populations, reflecting an interplay of river proximity, philopatry and resource availability. The stronger philopatry and constrained range of *A. fallax* imply heightened vulnerability to localized bycatch pressure near natal rivers, whereas *A. alosa*’s broader dispersal and higher origin diversity expose multiple populations to risk at a regional scale. These species-specific dispersal capacities and metapopulation structures critically shape bycatch vulnerability. Incorporating natal-origin and dispersal data into transnational, ecosystem-based management—such as targeted temporal or spatial fishing restrictions at mixing hotspots—will be essential to safeguard metapopulation dynamics, mitigate bycatch mortality, and maintain ecological connectivity among European shad populations.

## Introduction

Bycatch, the incidental capture of non-target species in marine commercial and artisanal fisheries, is a major threat to marine ecosystems (Davies *et al*., 2009; Gray and Kennelly, 2018). Depending on its fate, bycatch can be classified as landed bycatch (retained for consumption or sale) or discards (returned to the sea, alive or dead; Gray and Kennelly, 2018). This issue affects a wide range of marine fauna, including birds, turtles, marine mammals, and fish species, and poses a considerable challenge to conservation efforts (Barreiro *et al*., 2025; Bethoney *et al*., 2017; Chuenpagdee *et al*., 2003). In addition, effects of bycatch may interact with the ongoing climate change, creating new ecological interactions and compounding existing anthropogenic threats (Hazen *et al*., 2018; Poloczanska *et al*., 2013). Among the species affected by bycatch, anadromous fish— those migrating from marine habitats to rivers to spawn (McDowall, 1988, 2001)—are particularly of relevance, as many are classified as protected, endangered or threatened (PETS; ICES, 2022) and are experiencing global population declines (Limburg and Waldman, 2009). These species play a key role in maintaining ecosystem connectivity by transporting marine-derived nutrients to freshwater habitats, supporting nutrient cycling and food web dynamics (Hall *et al*., 2012; Poulet *et al*., 2022). Their depletion disrupts vital ecological processes/cycles, jeopardizing and leading to the loss of ecosystem services that benefit both local human communities and the broader environment across territories and administrative boundaries (Almeida *et al*., 2023; Ashley *et al*., 2023; Hall *et al*., 2012; Limburg and Waldman, 2009; Poulet *et al*., 2022). Beyond their role in ecosystem functioning, the anadromous clupeids allis shad *Alosa alosa* (L. 1758) and twaite shad *Alosa fallax* (Lacépède 1803), collectively known as European shads, are ecologically and socio-economically significant. Historically, they have supported inland commercial and recreational fisheries, providing provisioning services such as protein supply, cultural heritage, and traditional fisheries embedded in local folklore (Almeida *et al*., 2023; Ashley *et al*., 2023). However, their populations have declined dramatically, raising concerns about their conservation and the sustainability of related human activities.

*Alosa alosa* and *A. fallax*, once widespread from Scandinavia to Morocco and the Mediterranean Sea, have experienced significant declines in both distribution range and abundance along the Atlantic coast (Baglinière, 2000). Consequently, the sustainability of their inland commercial and recreational fisheries came into question, leading to increasing academic and fisheries management interest in their conservation (Braga *et al*., 2022; Elie *et al*., 2000; Mota *et al*., 2015; Rougier *et al*., 2012; Stratoudakis *et al*., 2016). Notably, the collapse of major European populations, such as those in the Gironde-Garonne-Dordogne system in France (Rougier *et al*., 2012) and the Minho River system on the Spain-Portugal border (Mota *et al*., 2015), underscores the severity of their decline. These declines are attributed to cumulative human impacts throughout their life cycle. While anthropogenic impacts in freshwater habitats are well-documented (Aprahamian *et al*., 2003; Limburg and Waldman, 2009; Taverny *et al*., 2000a), the effects of anthropogenic mortality at sea, where *A. alosa* and *A. fallax* migrate to feed and grow for several years, remain underexplored (Aprahamian *et al*., 2010; Davies *et al*., 2020; King and Roche, 2008; Nachón *et al*., 2016; Trancart *et al*., 2014). The declining status of European shad populations necessitates a thorough understanding of their specific threats. Both *Alosa* species exhibit vulnerability to bycatch due to their schooling behaviour, especially when aggregating to begin the spawning migrations and reliance on coastal/estuarine habitats that extensively overlap with fishing operations (King and Roche, 2008; Maitland and Lyle, 2005; OSPAR, 2009). Consequently, European shad bycatch has been widely documented across their distribution range over multiple decades (ICES, 2014; King and Roche, 2008; La Mesa *et al*., 2015; Nachón *et al*., 2016; Sabatié, 1993; Trancart *et al*., 2014). The resulting loss of pre-reproductive individuals may critically impair juvenile recruitment, further exacerbating population declines, a particularly severe consequence for the semelparous *A. alosa* (King and Roche, 2008). Nevertheless, comprehensive assessments of bycatch magnitude and demographic impacts remain scarce, hampered by inconsistent reporting protocols across EU Member States and persistent species misidentification (Baglinière *et al*., 2003; DiadSea Interreg Atlantic Area project; Nachón *et al*., 2016; OSPAR, 2009).

In this context, the mixing of shad populations at sea and in coastal habitats remains to be known (Nachón *et al*., 2020). Shad populations are defined by their native rivers, where adults spawn and juveniles migrate during their first year of life to marine habitats to feed and grow for several years (Aprahamian *et al*., 2003; Baglinière *et al*., 2003; Taverny and Elie, 2001). Upon reaching maturity, individuals return to their natal rivers to reproduce, maintaining genetic and ecological connectivity across generations, although a proportion stray into non-natal rivers (more frequently between neighbouring river basins), facilitating gene flow and demographic exchange between populations (Alexandrino *et al*., 2006; Jolly *et al*., 2012; Martin *et al*., 2015; Rougemont *et al*., 2022; Sabatino *et al*., 2022). During the marine phase, some individuals remain near the mouths of their natal rivers, while others disperse hundreds of kilometres away alongside the coast (Nachón *et al*., 2020). This variability in dispersal behaviour highlights the complexity of population mixing at sea and raises questions about its implications for connectivity and conservation. Such heterogeneity in movement prompts the use of intrinsic ‘natural tags,’ notably otoliths—calcified ear stones in teleost fishes—to resolve fine-scale connectivity. Certain trace elements and isotopes accrete continuously in otolith growth layers in proportion to ambient water chemistry, especially in freshwater environments (Campana and Thorrold, 2001; Daverat *et al*., 2016; Walther and Thorrold, 2006). Because these metabolically inert structures retain distinct natal-water signatures in their cores, they provide a retrospective record of both natal origin and subsequent dispersal patterns (Campana and Thorrold, 2001; Daverat *et al*., 2011; Martin *et al*., 2013a, b; Nachón, 2017; Walther *et al*., 2008). Otolith microchemistry —through Sr/Ca, Ba/Ca and ^87^Sr/^86^Sr ratios as natal tracers—has proven useful for tracing the natal origins of individuals in *A. alosa* and *A. fallax* (Martin *et al*., 2015; Nachón *et al*., 2020; Randon *et al*., 2018), offering a promising approach for examining population behaviour at sea and providing insights into the impact of bycatch on these species. In this context, understanding the effects of bycatch on the connectivity and diversity of natal origins is critical for informing conservation strategies. Therefore, this study aims to: (1) investigate the population connectivity and natal origins of *A. alosa* and *A. fallax* using otolith microchemistry from bycaught individuals; (2) infer whether bycatch impacts differ among populations; and (3) discuss the conservation implications and outline future directions for management strategies.

## Materials and Methods

### Origin of bycatch European shad samples

This study examines European shad specimens bycaught in small-scale coastal fisheries along the western Iberian coast and subsequently landed at fish markets located near the vessels’ home ports. As these fisheries generally operate close to their landing harbours (Nachón *et al*., 2016, 2022a, b), the recorded bycatch locations are assumed to reflect ecologically relevant marine habitats used by European shads (Nachón *et al*., 2022c, d). A total of 120 specimens were opportunistically acquired between February 2017 and March 2021 (Table 1), spanning from Spain’s Ártabro Gulf to Portugal’s Mondego estuary (Figueira da Foz; Figure 1).

**Figure 1.**
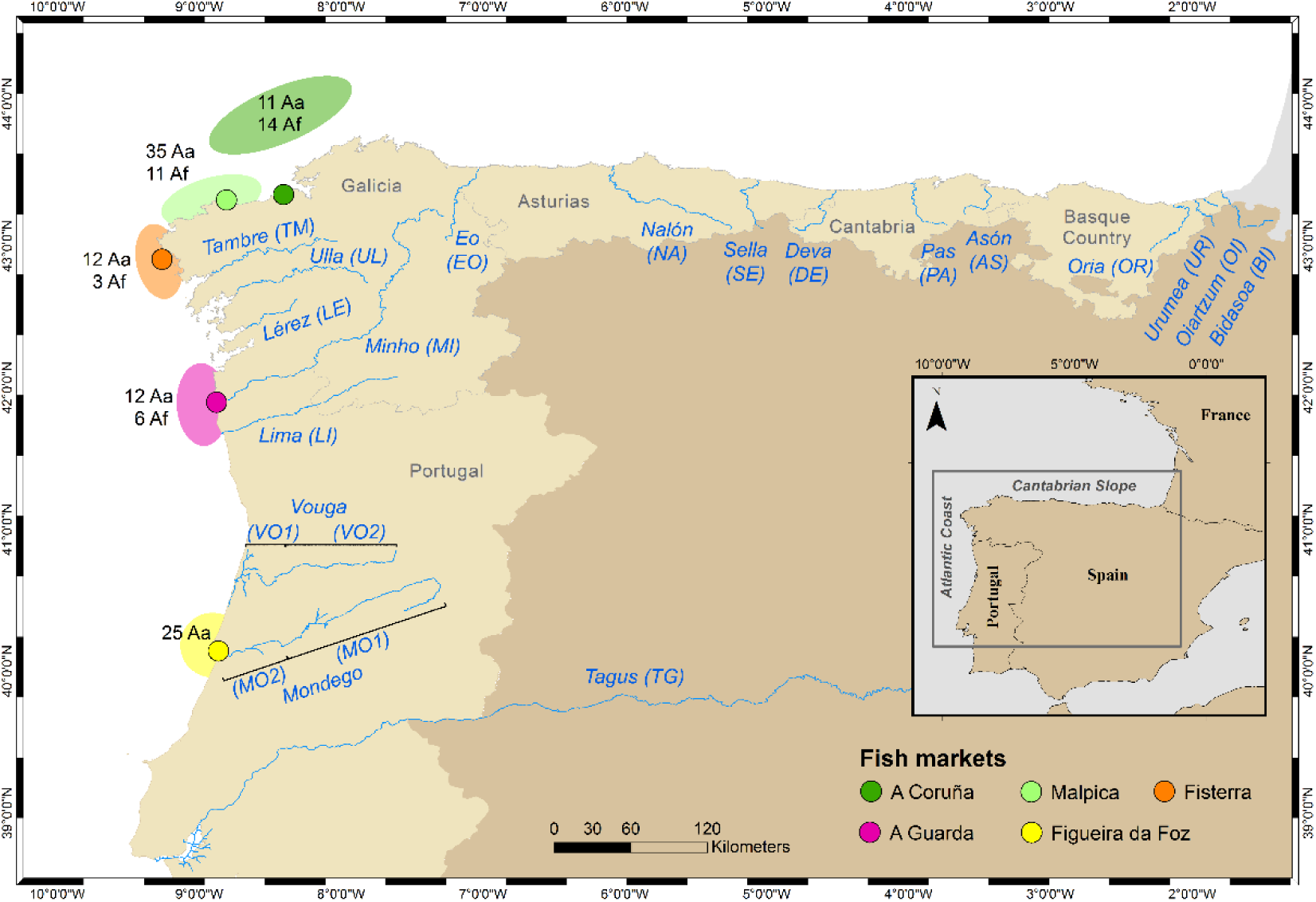
Study area along the western Iberian coast (northeastern North Atlantic), showing fish-market landing sites where bycatch specimens of Allis shad *Alosa alosa* (L. 1758) and Twaite shad *Alosa fallax* (Lacépède 1803) were obtained. Ellipses coloured by market indicate the approximate coastal capture locations, with sample counts for each species (Aa = *Alosa alosa*; Af = *Alosa fallax*). Major rivers on the Atlantic and Cantabrian coasts of the Iberian Peninsula where these species have been reported (Barber-O’Malley *et al*., 2022) are shown as potential natal sources and were included in the reference database for origin assignments. The Vouga and Mondego rivers are each divided into two segments (1 & 2) to reflect contrasting geologies and Sr-isotope signatures rather than averaging across the entire watershed.

**Table 1.** Number of bycaught individuals and biometric values (mean ± S.D.) of Allis shad *Alosa alosa* (L. 1758) and Twaite shad *Alosa fallax* (Lacépède 1803) recorded at fish markets, ordered latitudinally from north to south along the northwest Iberian Peninsula, with corresponding sampling dates (months and years).

| Fish market | Dates (months/years) | *A. alosa* |  |  |  | *A. fallax* |  |  |  |
| --- | --- | --- | --- | --- | --- | --- | --- | --- | --- |
|  |  | *N* | *L<sub>T</sub>* (cm) | *M* (g) | Gill rakers (*n*) | *N* | *L<sub>T</sub>* (cm) | *M* (g) | Gill rakers (*n*) |
| A Coruña | Ja-Fe-Mar/2021 | 11 | 54.6±8 | 1690.3±814.6 | 130.7±7.6 | 14 | 43.6±2.1 | 690.5±97.5 | 45.6±2.5 |
| Malpica | Dec/2019; Oct/2020; Fe-Mar/2021 | 31 | 51.7±3.8 | 1404.7±386 | 128.5±6.6 | 11 | 44.4±3.3 | 776.2±203.4 | 48.6±4.6 |
| Fisterra | Ja-Fe/2020 | 10 | 54±6.4 | 1843±403.6 | 132.9±6.3 | | | | |
| A Guarda | Ja-Fe-Mar/2021 | 12 | 49.8±4.5 | 1161±432.8 | 135.6±9.9 | 6 | 44.9±5.8 | 834.2±308.2 | 49.7±3.9 |
| Figueira da Foz | Fe-Mar/2017 | 25 | 60.6±4.9 | 2378.8±642.3 | 125.5±12.2 | | | | |
| Total | | 89 | 54.5±6.5 | 1730.5±692.4 | 129.5±9.3 | 31 | 44.2±3.4 | 748.7±192.5 | 47.5±4 |
| Abbreviations: S.D., standard deviation; Ja, January; Fe, February; Mar, March; Oct, October; Dec, December; *N*, number of samples; *L<sub>T</sub>*, total length; *M*, mass; *n*, count. |  |  |  |  |  |  |  |  |  |

In the laboratory, species identification was conducted through gill raker counts on the first branchial arch following Alexandrino *et al*. (2006), confirming 89 *A. alosa* and 31 *A. fallax* individuals. Most specimens (*n*=95: 64 *A. alosa* and 31 *A. fallax*) represented bycatch from Galician inshore gillnet fisheries targeting commercial species—e.g., European sea bass *Dicentrarchus labrax* (L. 1758), European hake *Merluccius merluccius* (L. 1758), acquired through first-sale records at four fish markets A Coruña, Malpica, Fisterra, and A Guarda (Table 1 and Figure 1). Fisheries context can be found in Nachón *et al*. (2016). The remaining 25 *A. alosa* individuals were obtained from specimens caught as bycatch in commercial fixed gillnets operating near Figueira da Foz (Mondego estuary, Portugal) during February-March 2017 (Table 1 and Figure 1). Otoliths were extracted from the cephalic regions of bycaught specimens following the protocol described in Martin *et al*. (2015). These otoliths were then processed to reconstruct dispersal trajectories and natal origins, providing valuable biological data for bycatch impact assessment while preventing further exploitation of these vulnerable species—an approach that addresses key ethical concerns in threatened species research.

### Natal origin baseline: water and juvenile tracers

In this study, we expanded the microchemical datasets for French and Portuguese rivers, originally analysed by Randon *et al*. (2018), by adding newly collected water samples and juvenile otoliths (Table 2). Because elemental uptake from water into otoliths is mediated by physiological and environmental processes (Campana, 1999; Campana and Thorrold, 2001; Kalish, 1989), we need to calibrate this water–otolith relationship using paired water and juvenile-otolith samples. Such calibration is crucial for predicting otolith signatures in rivers where juvenile sampling is impractical—both due to challenges of early-life sampling and the species’ sensitive conservation status.

**Table 2.**
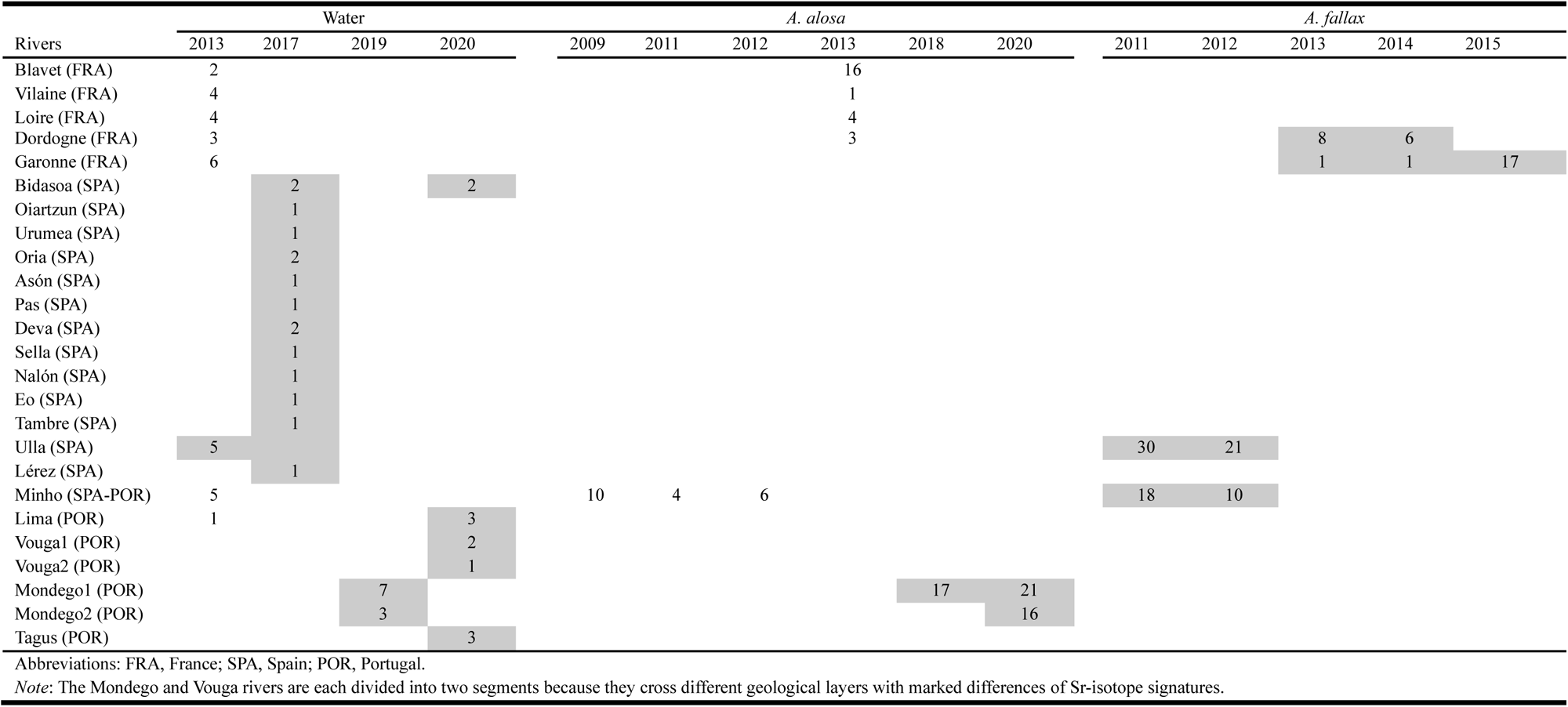
Number of water samples and juvenile otoliths used to build the natal-origin baseline for Allis shad *Alosa alosa* (L. 1758) and Twaite shad *Alosa fallax* (Lacépède 1803). Water samples and juvenile otoliths of *A. alosa* from French and Portuguese rivers (Randon *et al*., 2018) were supplemented by new water samples and juvenile otoliths of both species from rivers along the Cantabrian slope and Atlantic coast rivers of northwestern Iberian Peninsula. Rivers are listed from north to south; values indicate sample

The original calibrations from Randon *et al*. (2018)—focused on *A. alosa* juveniles— were reinforced with fresh water and otolith collections from the Mondego River (Portugal, Table 2), where we sampled upstream and downstream of a rehabilitated fish-pass dam to evaluate connectivity-mediated recruitment success (see Belo *et al*., 2021 for rehabilitation details). Juveniles were captured following Martin *et al*. (2015), using purse-seine nets in upper-estuarine and freshwater reaches during seaward migration, ensuring natal origin confirmation prior to marine entry (Walther *et al*., 2008). To extend the calibration to *A. fallax*, we incorporated archived juvenile otoliths from the Ulla and Minho rivers (Galicia, Spain)—originally sampled in Nachón *et al*. (2016) and housed in the University of Santiago de Compostela (USC) collection (Table 2). We also added archived *A. fallax* juvenile otoliths from systematic sampling programs in the Garonne and Dordogne basins (Table 2), stored at the National Institute for Agricultural Research and the Environment (INRAE) research institute in Cestas (Bordeaux, France).

Regarding water chemistry, we reinforced reference coverage from Randon *et al*. (2018) and extended our spatial scope to the Cantabrian-slope rivers in Spain and to the Vouga, Mondego, and Tagus systems in Portugal (Table 2). Because the Vouga and Mondego systems traverse contrasting geologies with markedly different Sr-isotope signatures, we divided each into two segments (Mondego 1 & 2; Vouga 1 & 2; see Figure 1) rather than averaging across the entire watershed. For the Mondego River specifically, we sampled immediately upstream and downstream of its first rehabilitated barrier—thereby linking river-segment water chemistry to juvenile-otolith microchemistry in each reach. Water samples were collected between May and September—spanning the spawning and post-spawning seasons—following Martin *et al*. (2015), and targeted areas adjacent to known or potential spawning grounds.

Although our calibration incorporated French rivers, natal-origin assignments of bycaught adults were confined to the Iberian Peninsula (Figure 1). This spatially explicit framework spanned 560 km of the Atlantic façade (Tagus River to Ártabro Gulf) and another 560 km along the Cantabrian coast (Ártabro Gulf to Bidasoa River at the Spain-France border), thereby encompassing both core regional distribution areas and typical dispersal ranges of both species (Jolly *et al*., 2012; Martin *et al*., 2015; Mota *et al*., 2016; Nachón *et al*., 2019a, b). This spatial framework is specifically tailored to assign adults accidentally caught in the coastal waters of northwestern Iberia to their natal rivers.

### Ethical Statement

European shad bycaught specimens acquired from fish markets were already deceased upon landing, eliminating any need for live animal experimentation. By treating otoliths as post-mortem “black-box” records, we extract maximal life-history information to reveal the impacts of bycatch on threatened European shad populations.

*Alosa alosa* juveniles from the Mondego River (Portugal) were captured during scientific sampling campaigns under authorization from the Instituto da Conservação da Natureza e das Florestas (ICNF; Licença 226/2020/CAPT). Sampling was conducted with rigorous attention to fish welfare and restricted to the minimum number of specimens necessary to achieve the study’s objectives. Otolith analyses were focused on individuals collected upstream and downstream of a rehabilitated dam to assess both habitat reconnection and bycatch impacts. To further reduce live captures, we incorporated existing juvenile-otolith microchemistry datasets for *A. alosa* and archived *A. fallax* otoliths. We also employed statistical models to generate “virtual” juveniles for under-sampled rivers. Finally, all otolith microchemistry data directly inform targeted conservation measures aimed at minimizing bycatch mortality and strengthening population resilience.

### Water and otolith microchemistry analysis

As established previously in studies by Martin *et al*. (2015), Nachón *et al*. (2020), and Randon *et al*. (2018), the elements Sr and Ba (in relation to Ca) and the isotopic ratios (^87^Sr/^86^Sr) were selected as key markers for determining the natal origin of European shads. Elemental concentrations of Ca, Sr, and Ba in both water and otoliths were analysed using solution-based Inductively Coupled Plasma Mass Spectrometry (ICP-MS). Isotopic ratios (^87^Sr/^86^Sr) were determined using a Nu-Plasma Multi-Collector ICP-MS. Otolith material was extracted with a femtosecond laser, tracing two C-shaped specular trajectories 40 microns from the core and 60 microns wide. Detailed protocols for water and otolith analysis are described in Martin *et al*. (2013a) and Martin *et al*. (2015), respectively. Regarding water samples from the Lima, Vouga, Mondego, and Tagus rivers, collected in 2019 and 2020, they were analysed by the Central Analysis Laboratory of Aveiro University. The same standards and methodologies were applied as in previous studies, ensuring comparison and consistency. Sr, Ca, and Ba were quantified using ICP-OES (Jobin Yvon Activa M), following the ISO 11885 standard, while isotopic ratios (^87^Sr/^86^Sr) were determined using ICP-MS (Thermo X Series).

### Natal origin assignment model for European shad bycatch at sea

Natal origin assignment for shads caught at sea involved three main steps: (1) developing a regression model for tracer concentrations in juvenile otoliths, (2) selecting, training, and validating the most effective classification model, and (3) calculating the probabilities of natal origin for each adult individual of both species. These steps are detailed below.

#### 1. Regression model for tracer concentrations in juvenile otoliths

We developed a model relating water chemistry (Sr/Ca, Ba/Ca, and ^87^Sr/^86^Sr isotope ratios) to juvenile otolith signatures using data from rivers where both water samples and juvenile specimens were available. The ^87^Sr/^86^Sr isotope ratios in otoliths ratios proved particularly valuable for natal origin assignment, as they precisely reflect stream water values without trophic fractionation (Kennedy *et al*., 2000; Martin *et al*., 2013a), providing unambiguous local-scale discrimination of European shad populations (Martin *et al*., 2015; Nachón *et al*., 2020; Randon *et al*., 2018).

For Sr and Ba concentrations, we applied linear regression models based on established water-otolith relationships (Martin *et al*., 2013a; Walther and Thorrold, 2008). Model validation was performed using Sr/Ca and Ba/Ca ratios in a synthetic dataset, incorporating data from Randon *et al*. (2018). For Sr/Ca, regressions were highly significant in both *A. alosa* (F-test; *F* = 44.96, d.f. = 5, *P*-value = 0.001, R² = 0.88) and *A. fallax* (F-test; *F* = 21.68, d.f. = 2, *P*-value = 0.042, R² = 0.88). For Ba/Ca, regression models explained biologically meaningful proportion of variance (*A. alosa*: F-test; *F* = 5.67, d.f. = 5, *P*-value = 0.063, R² = 0.44; *A. fallax*: F-test; *F* = 12.58, d.f. = 2, *P*-value = 0.072, R² = 0.79), though these relationships were marginally non-significant at α = 0.05. This pattern likely reflects both the high natural variability of barium in otoliths (Bareille *et al*., 2024; Martin *et al*., 2013a; Walther *et al*, 2008) and limited sample sizes for some rivers. Despite the lack of statistical significance, we retained Ba/Ca models because: (1) Barium provides complementary spatial information to strontium (Bareille *et al*., 2024; Martin *et al*., 2013a; Walther *et al*. 2008), (2) The R² values (0.44-0.79) indicate biologically meaningful effects (Cohen, 1988; Nakagawa and Cuthill, 2007), and (3) multi-tracer integration increases assignment accuracy (Martin *et al*., 2013a; Walther *et al*. 2008). While Ba/Ca alone shows limited discriminatory power, its combined use with other tracers reduces misclassification risks in variable environments. The variance structure of Ba/Ca was therefore incorporated into synthetic datasets to maintain methodological consistency across all tracers. These models generated the reference data needed for natal origin assignment in marine-bycaught shads. For rivers with only water chemistry data (Table 2), we created synthetic datasets of 60 juvenile otoliths per river, assuming a normal distribution (X ∼ N(µi, σi)) where µi (mean) and σi (standard deviation) were derived from rivers with paired water-otolith measurements (Table 2). Model parameters, calculated as sample estimates of mean and standard deviation, were derived exclusively from rivers where both water chemistry data and juvenile otolith samples were available (Table 2). When available juvenile otoliths numbered fewer than 60 specimens per river, we augmented biological samples with simulated data using identical protocols to ensure methodological uniformity. Dissimilarities in microchemical otolith composition among different rivers were then quantified using Canberra distance (Lance and Williams, 1966, 1967).

#### 2. Model selection, training and validation

An exploratory analysis was conducted to compare the performance of four classification algorithms (Naïve Bayes, Quadratic Discriminant Analysis—QDA, Random Forest, and Stagewise Additive Modelling with a Multi-class Exponential loss function—SAMME) for assigning natal origin to European shad bycatch adults. As no significant differences were found in average classification accuracy, QDA was selected due to its effectiveness and lower computational cost. QDA is particularly suitable for classification tasks with moderate sample sizes and has been shown to perform well while maintaining relatively low computational requirements (e.g., Friedman, 1989; Hastie *et al*., 2009; Smoliński *et al*., 2020).

The model was trained using 70% of available juvenile samples, with the remaining 30% reserved for validation. To optimise performance, cross-validation with repeated subsampling was applied on the training dataset, enabling selection of the optimal hyperparameters. Specifically, five repetitions of 10-fold cross-validation were performed to ensure robust model selection and mitigate overfitting, as recommended by Hastie *et al*. (2009). This cross-validation approach was applied to the training dataset, while the test dataset (comprising the remaining 30% of the samples) was used to estimate accuracy and assess the model’s practical performance.

#### 3. Bycatch European shads’ natal rivers allocation

After adjusting and validating the classification model, the probabilities of natal origin for each adult individual of both species were calculated. We established three probability thresholds (80%, 65%, and 50%) to assess prediction confidence across a range of reliability. Adults meeting or exceeding a threshold were assigned to the corresponding natal river, while specimens with probabilities below the 50% threshold were classified as unknown due to insufficient assignment confidence. The use of multiple thresholds allows us to evaluate the robustness of the model and identify specimens with lower assignment confidence, which could provide insights into potential regional patterns in misclassification. To characterise the spatial movements of bycaught individuals, displacement direction (latitudinal or longitudinal) and distance from the inferred natal river to the approximate bycatch location were classified. Straight-line distances were estimated from the river mouth to the midpoint of the corresponding fish market’s bycatch locations. Distance classes followed the categories proposed by Martin *et al*. (2015), with the addition of an ultra-short range (<20 km) to capture finer-scale movements.

All analyses were conducted using R version 4.4.3 (R Core Team, 2025). Full computer codes and data necessary to replicate the analyses and produce the results reported in the manuscript are available in the otoshad repository (Anonymous, 2025).

## Results

### Natal origin baseline: water and juvenile tracers

Canberra distance analysis of strontium (Sr/Ca), barium (Ba/Ca), and the ^87^Sr/^86^Sr isotope ratio revealed significant microchemical differentiation between juveniles from different river systems (Figure 2). Canberra distance analysis demonstrated systematic microchemical differentiation across spatial scales.

**Figure 2.**
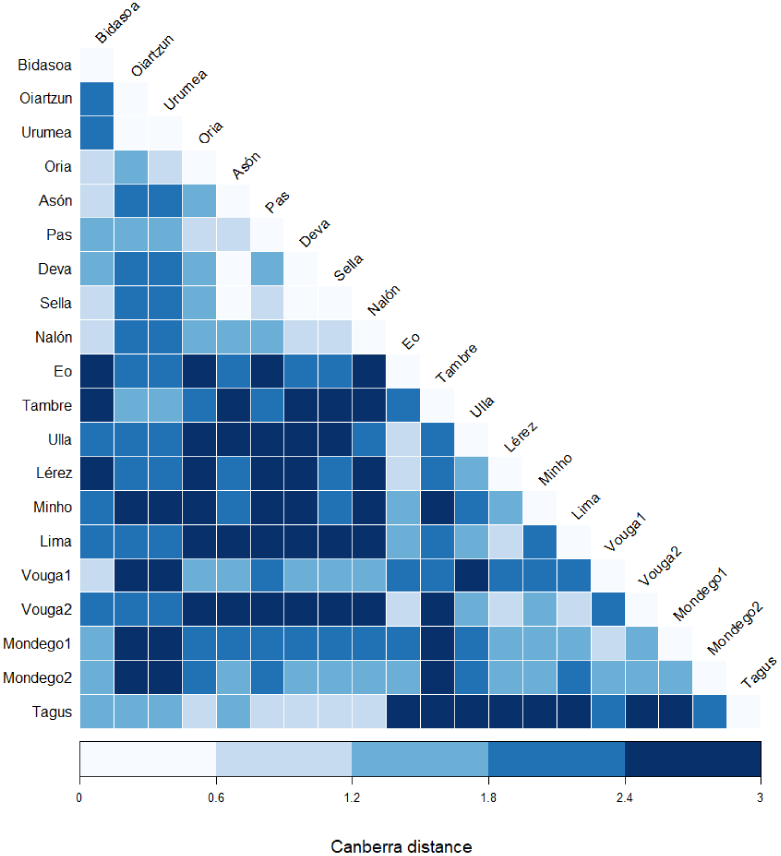
Canberra distance matrix among river systems based on combined juvenile otolith microchemical signatures (Sr/Ca, Ba/Ca and ⁸⁷Sr/⁸⁶Sr) from Allis shad *Alosa alosa* (L. 1758) and Twaite shad *Alosa fallax* (Lacépède 1803). Rivers are ordered geographically from Cantabrian to Atlantic systems along both axes, with Mondego and Vouga rivers split into two segments to reflect contrasting ⁸⁷Sr/⁸⁶Sr signatures. Cell colours indicate the magnitude of Canberra distance according to the colour scale included within the figure.

At the regional level, granitic rivers of the Atlantic coast (Tambre to Tagus) showed pronounced divergence from calcareous systems of the Cantabrian slope (Bidasoa to Eo), with 71% of inter-group comparisons exhibiting distances >1.8 (40% at 2.4–3.0; 31% at 1.8–2.4). The Tagus River constituted a striking exception—its sedimentary geology produced signatures akin to Cantabrian calcareous rivers (Pas, Deva, Sella, Nalón), with distances of 0.6–1.2 (9% of comparisons). While these values reflect closer geochemical affinity than typical inter-group pairs, they remain diagnostically distinguishable. At the individual river scale, 81.25% of pairwise comparisons showed strong differentiation (distances >1.2), with 61.25% falling within the 1.8–3.0 range. Intermediate distances (1.2–1.8) accounted for 24.6% of cases, while only 14.13% exhibited low differentiation (0.6–1.2). Crucially, the minimal-distance category (0–0.6; 2.09% of comparisons) was restricted to immediately adjacent systems (e.g., Oiartzun-Urumea, Deva-Sella), where microchemical overlap poses negligible ecological consequences. This systematic variation confirms that bedrock geology, specifically the differentiation between granite and sedimentary substrates, rather than geographic proximity, primarily drives otolith microchemical fingerprints. However, within these broad geological categories, smaller-scale variability likely arises from the heterogeneity of substrates within different catchment areas.

The differentiation was consistent across species, with similar microchemical patterns observed in juvenile *A. alosa* (Figures 3a-b-c) and *A. fallax* (Figures 4a-b-c). The ^87^Sr/^86^Sr isotope ratio emerged as the most effective river discriminator, revealing a clear biogeochemical gradient: Cantabrian slope rivers consistently showed values <0.710, while Atlantic façade rivers typically exceeded 0.715 (with rare exceptions). Barium concentrations, while variable within individual rivers, enhanced discrimination when combined with strontium and its isotope ratios. Notably, Cantabrian slope rivers exhibited greater microchemical homogeneity compared to the Atlantic façade, reinforcing the geological basis of these regional fingerprints.

**Figure 3.**
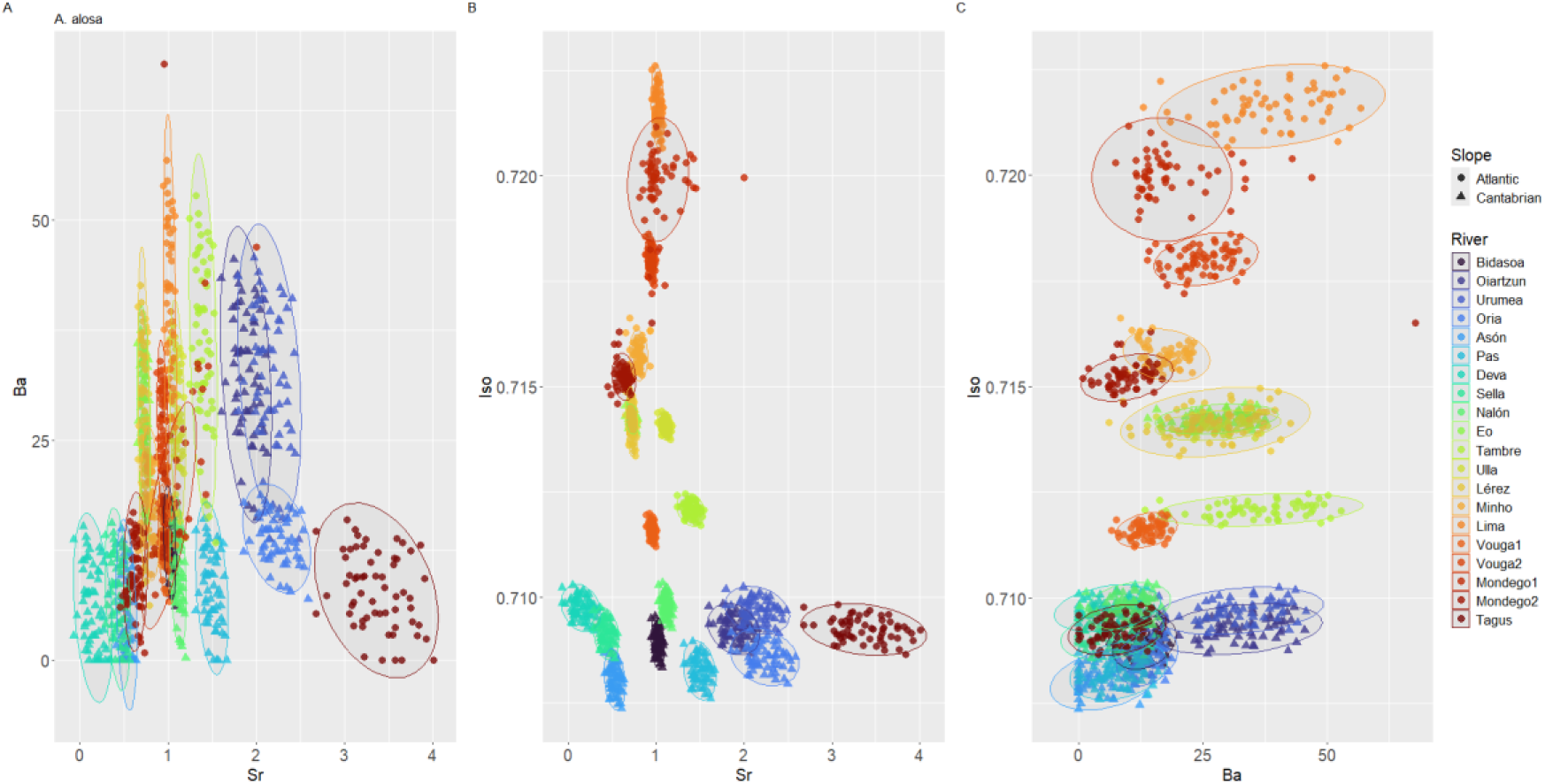
Differentiation of juvenile otolith microchemical signatures in *Alosa alosa* (L. 1758) across river systems based on Sr/Ca, Ba/Ca and ⁸⁷Sr/⁸⁶Sr. Panels show (a) Sr/Ca vs Ba/Ca, (b) ⁸⁷Sr/⁸⁶Sr vs Ba/Ca, and (c) ⁸⁷Sr/⁸⁶Sr vs Sr/Ca. Juveniles (real and simulated; n = 60 per river) are colour-coded by river of origin, ordered geographically from the Cantabrian to the Atlantic coast. Colours follow the scale included in the figure.

**Figure 4.**
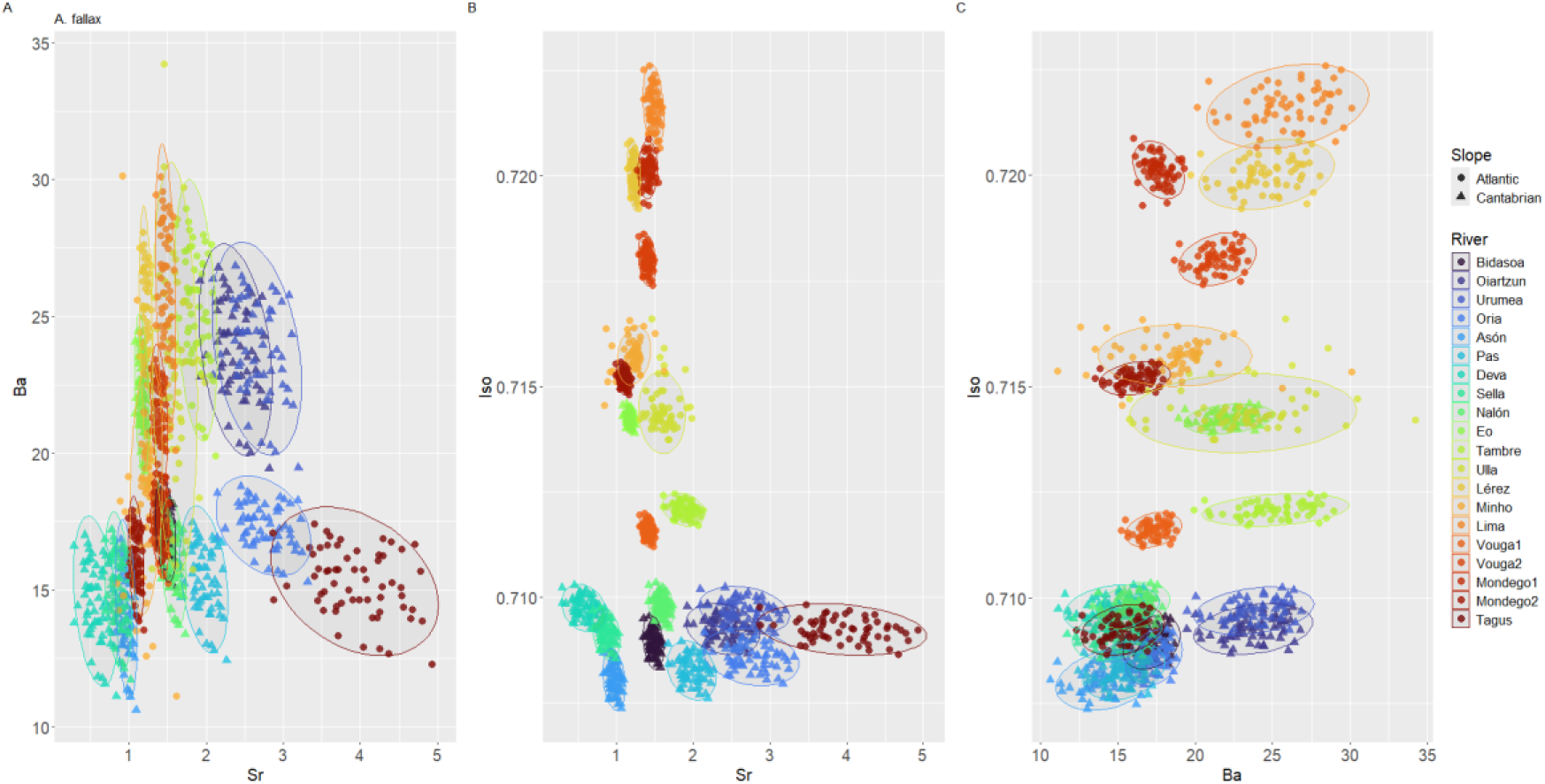
Differentiation of juvenile otolith microchemical signatures in Twaite shad *Alosa fallax* (Lacépede 1803) across river systems based on Sr/Ca, Ba/Ca and ⁸⁷Sr/⁸⁶Sr. Panels show (a) Sr/Ca vs Ba/Ca, (b) ⁸⁷Sr/⁸⁶Sr vs Ba/Ca, and (c) ⁸⁷Sr/⁸⁶Sr vs Sr/Ca. Juveniles (real and simulated; n = 60 per river) are colour-coded by river of origin, ordered geographically from the Cantabrian to the Atlantic coast. Colours follow the scale included in the figure.

### Model performance and bycatch European shads’ natal rivers allocation

The classification model demonstrated high predictive accuracy after 50 resampling iterations (5 repetitions of 10-fold cross-validation), ensuring robustness. In the training subset, classification accuracies were 94.76±0.02% for *A. alosa* (Figure 5a-b) and 96.83±0.01% for *A. fallax* (Figure 5c-d). These results were closely mirrored in the validation subset, with accuracies of 94.44% and 94.72%, respectively, indicating strong generalisation capacity. The consistency between training and validation results suggests that the model effectively captures microchemical variations among river systems while minimising overfitting.

**Figure 5.**
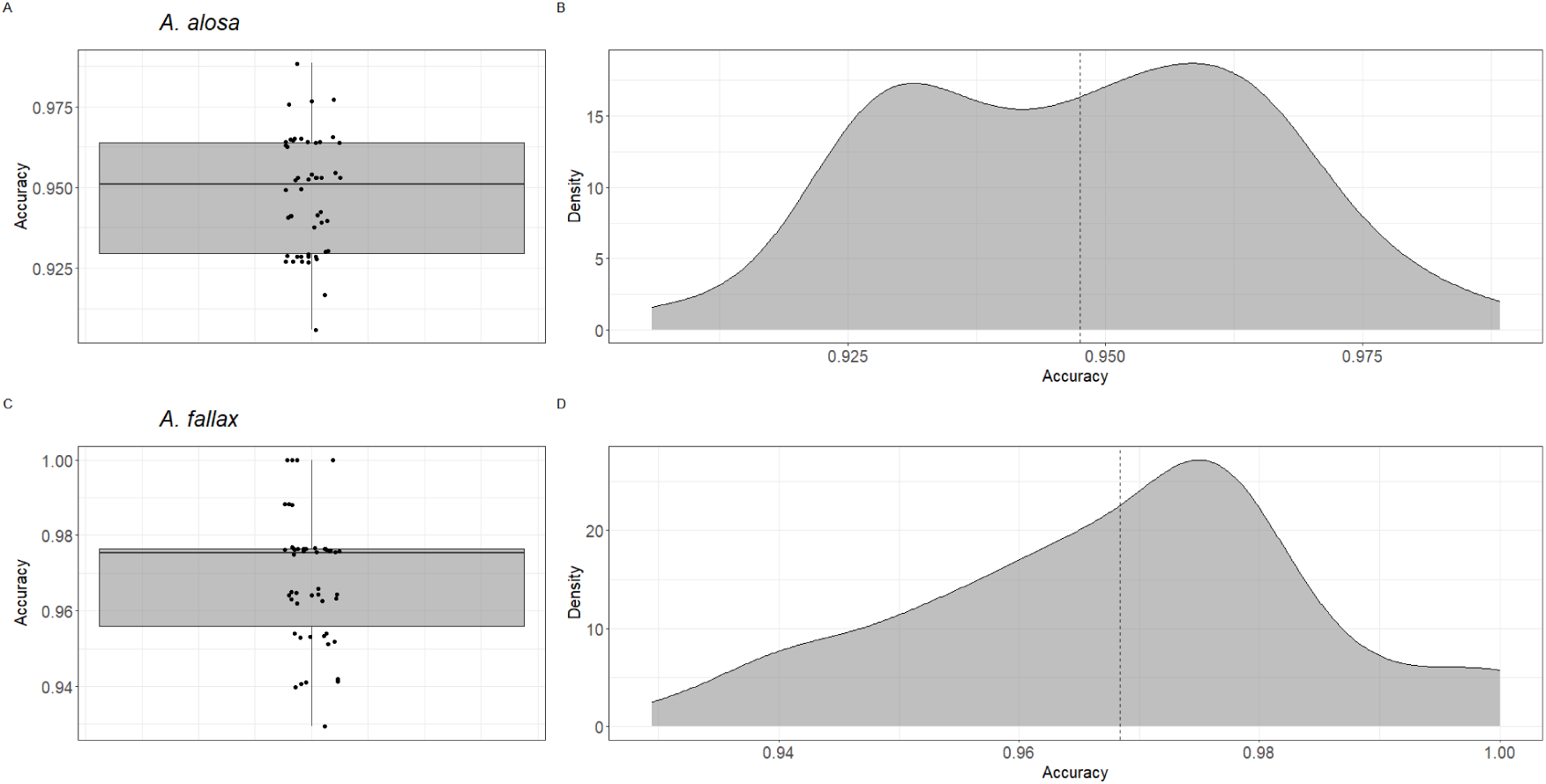
Classification model performance for juvenile Allis shad *Alosa alosa* (L. 1758) and Twaite shad *Alosa fallax* (Lacépede 1803) across 50 resamples using the training subset. Panels a and c show the classification accuracy achieved in each cross-validation iteration. Panels b and d display the corresponding accuracy distributions, with the dashed line indicating the mean accuracy.

All bycaught individuals were successfully assigned to their natal rivers, with probability thresholds of 50% for *A. alosa* (Figure 6a and Table 3) and 65% for *A. fallax* (Figure 6b and Table 4), achieving 100% assignment rates. The highest threshold (80%) confirmed high-confidence assignments for the vast majority of specimens: 88.76% of *A. alosa* (79/89) and 90.32% of *A. fallax* (28/31). This demonstrates robust model performance, with only a small fraction of specimens (11.6% and 8.8%, respectively) showing intermediate probability values (50-80%).

**Figure 6.**
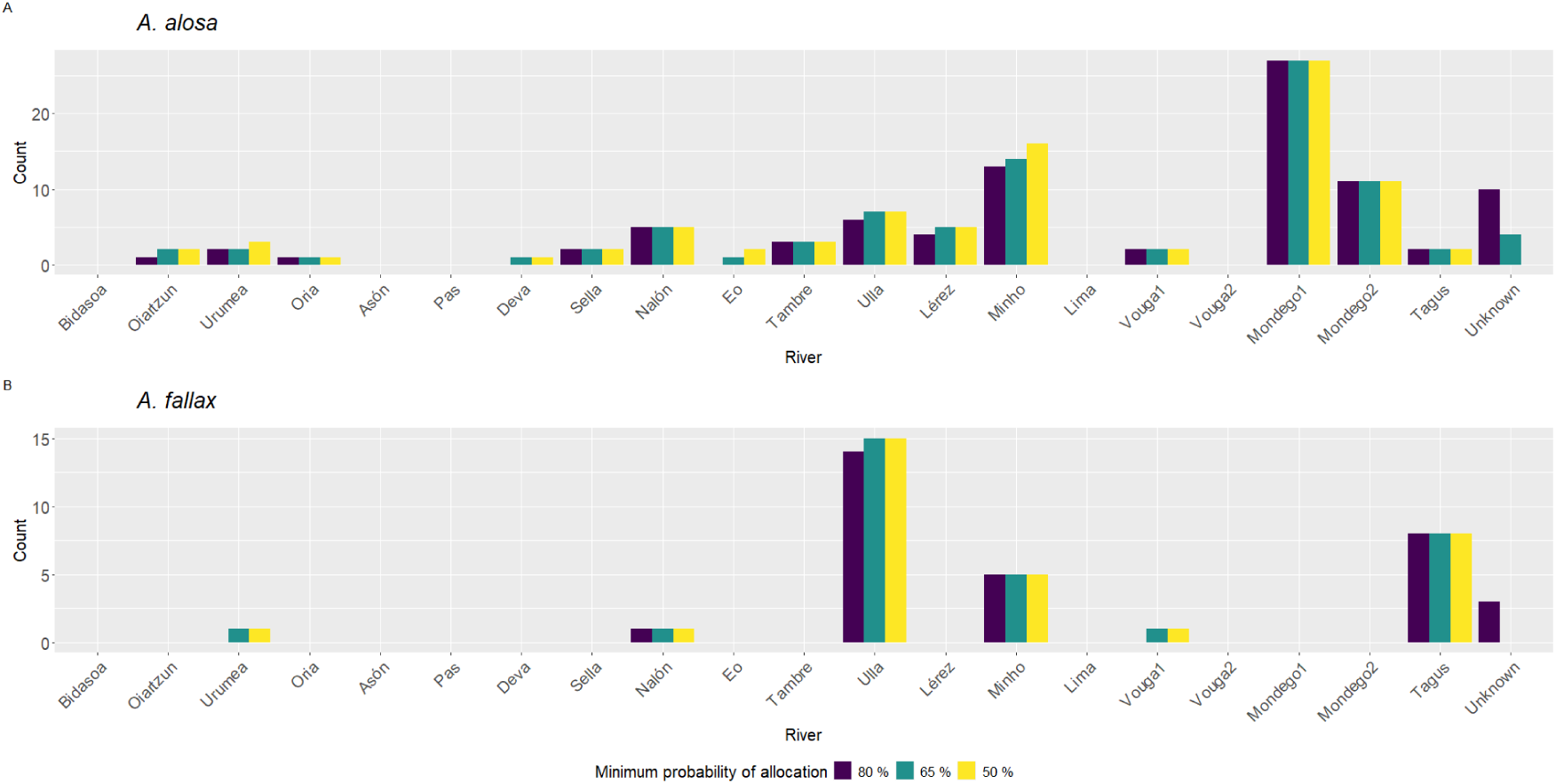
Number of Allis shad *Alosa alosa* (L. 1758) and Twaite shad *Alosa fallax* (Lacépede 1803) specimens assigned to their natal rivers at three minimum assignment-probability thresholds (80 %, 65 %, and 50 %). Bars show the number of individuals meeting each threshold; those below each threshold were classified as “Unknown”.

**Table 3.** Assignment of bycaught Allis shad *Alosa alosa* (L. 1758) specimens to their natal river (geographically ordered from north to south) at three minimum probability thresholds (80%, 65%, and 50%). Values are shown as number of assigned individuals (*N*) and percentage (%).

| Rivers | Probability thresholds |  |  |  |  |  |
| --- | --- | --- | --- | --- | --- | --- |
|  | 80% |  | 65% |  | 50% |  |
|  | *N* | % | *N* | % | *N* | % |
| Bidasoa | 0 | 0% | 0 | 0% | 0 | 0% |
| Oiartzun | 1 | 1.12% | 2 | 2.25% | 2 | 2.25% |
| Urumea | 2 | 2.25% | 2 | 2.25% | 3 | 3.37% |
| Oria | 1 | 1.12% | 1 | 1.12% | 1 | 1.12% |
| Asón | 0 | 0% | 0 | 0% | 0 | 0% |
| Pas | 0 | 0% | 0 | 0% | 0 | 0% |
| Deva | 0 | 0% | 1 | 1.12% | 1 | 1.12% |
| Sella | 2 | 2.25% | 2 | 2.25% | 2 | 2.25% |
| Nalón | 5 | 5.62% | 5 | 5.62% | 5 | 5.62% |
| Eo | 0 | 0% | 1 | 1.12% | 2 | 2.25% |
| Tambre | 3 | 3.37% | 3 | 3.37% | 3 | 3.37% |
| Ulla | 6 | 6.74% | 7 | 7.87% | 7 | 7.87% |
| Lérez | 4 | 4.49% | 5 | 5.62% | 5 | 5.62% |
| Minho | 13 | 14.61% | 14 | 15.73% | 16 | 17.98% |
| Lima | 0 | 0% | 0 | 0% | 0 | 0% |
| Vouga1 | 2 | 2.25% | 2 | 2.25% | 2 | 2.25% |
| Vouga2 | 0 | 0% | 0 | 0% | 0 | 0% |
| Mondego1 | 27 | 30.34% | 27 | 30.34% | 27 | 30.34% |
| Mondego2 | 11 | 12.36% | 11 | 12.36% | 11 | 12.36% |
| Tagus | 2 | 2.35% | 2 | 2.25% | 2 | 2.25% |
| Unknown | 10 | 11.24% | 4 | 4.49% | 0 | 0% |
| Total | 79 | 88.76% | 85 | 95.51% | 89 | 100% |

Abbreviations: *N*, number of individuals; %, percentage.
Note: “Unknown” comprises individuals whose maximum assignment probability fell below 50%. In the “Total” row, both *N* and % refer only to individuals correctly assigned at each probability threshold (i.e., “Unknown” individuals are excluded from these values).

**Table 4.**
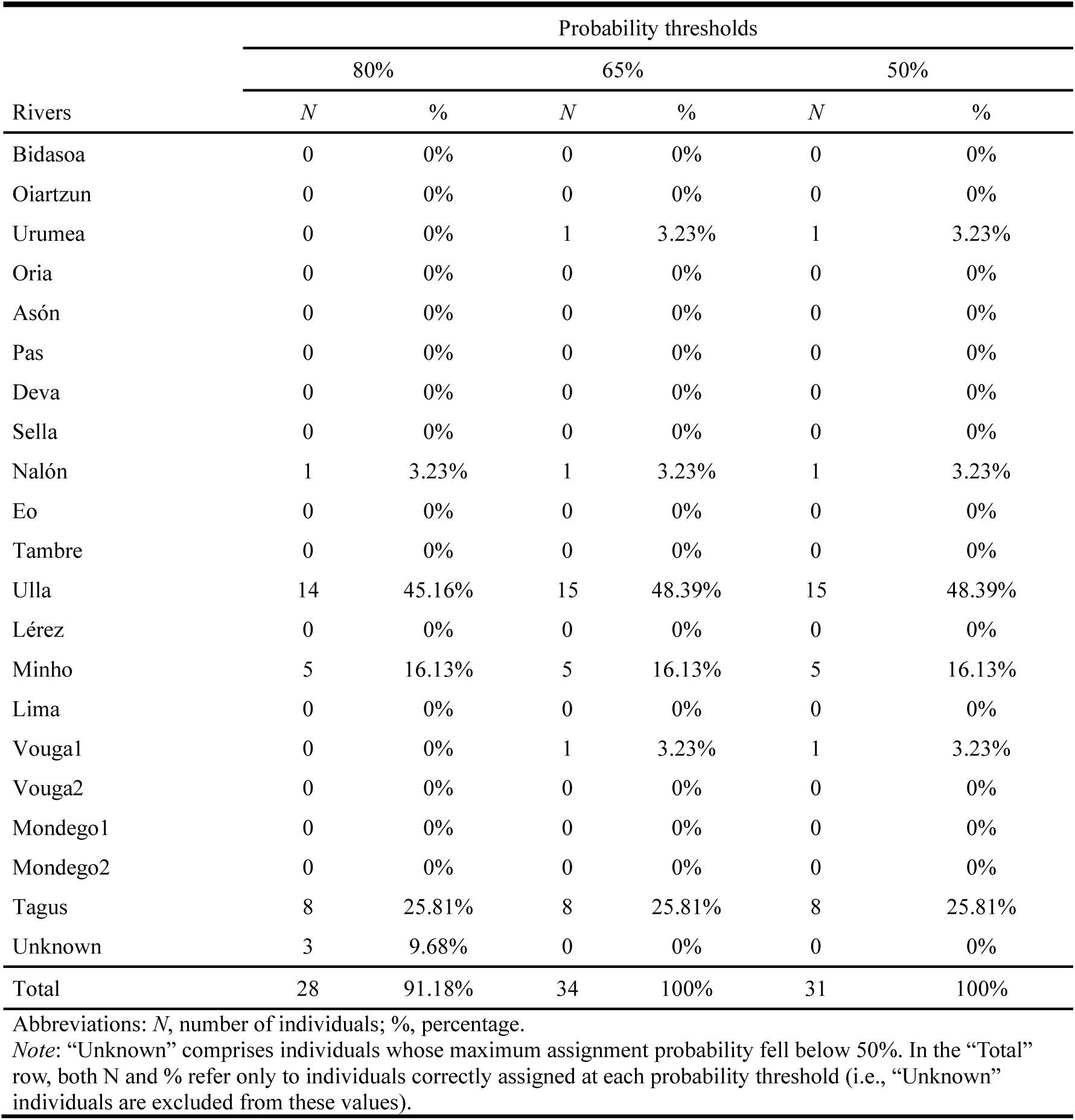
Assignment of bycaught Twaite shad *Alosa fallax* (Lacépède 1803) specimens to their natal river (geographically ordered from north to south) at three minimum probability thresholds (80%, 65%, and 50%). Values are shown as number of assigned individuals (*N*) and percentage (%).

Assignment results revealed contrasting patterns between species (Figures 7 and 8 and Tables 5 and 6). Natal-origin diversity in fish markets was significantly higher for *A. alosa* (mean = 7, range = 4-12) than for *A. fallax* (mean = 3, range = 2-3). For *A. alosa*, diversity varied substantially among markets: A Coruña, Fisterra and A Guarda showed the lowest diversity (4 origins each), Figueira da Foz intermediate diversity (9 origins), and Malpica the highest diversity (12 origins; Figure 7). The Mondego River represented both the most frequent and ubiquitous natal origin (present in all markets), followed by the Minho River (absent only from A Coruña). Most individuals exhibited latitudinal movements along the Atlantic coast, while longitudinal movements along the Cantabrian slope were less common (Table 7). Most displacements fell within the middle (100–300 km) and long (300–700 km) distance categories. Ultra-short movements (<20 km) were recorded only in the southern fish markets (A Guarda and Figueira da Foz), which are located at the mouths of the Minho and Mondego rivers, respectively—sites with well-established populations of these species. In contrast, individuals landed in the northern markets (e.g., Fisterra and Malpica) tended to originate from more distant rivers. In contrast, *A. fallax* exhibited limited variation, with only four natal origins identified: Nalón, Ulla, Minho, and Tagus rivers. In Malpica, only two of these origins were observed, compared to three in A Coruña and A Guarda (Figure 8). The Ulla River was the dominant natal origin in northern fish markets (A Coruña, Malpica), while the Minho River prevailed in the southern market (A Guarda). Interestingly, although the Tagus River is located south of the fish markets, its signal appeared only in the northernmost markets. Conversely, the Nalón River, situated to the northeast, was detected exclusively in the southernmost market. Spatial displacement patterns were more constrained than in *A. alosa*, reflecting the lower natal-origin diversity observed across fish markets. Most individuals exhibited latitudinal movements along the Atlantic coast, while longitudinal movements along the Cantabrian slope were rare and limited to a single individual in A Guarda (Table 8). Middle-distance displacements (100–300 km) predominated in the northern markets of A Coruña and Malpica, accounting for over 60% of cases in both. In contrast, bycatch in A Guarda showed a more heterogeneous pattern: 60% of individuals displayed ultra-short movements (<20 km), corresponding to the proximity of the Minho River population, while the remaining individuals were split between short (20–100 km) and long-distance (300–700 km) categories.

**Figure 7.**
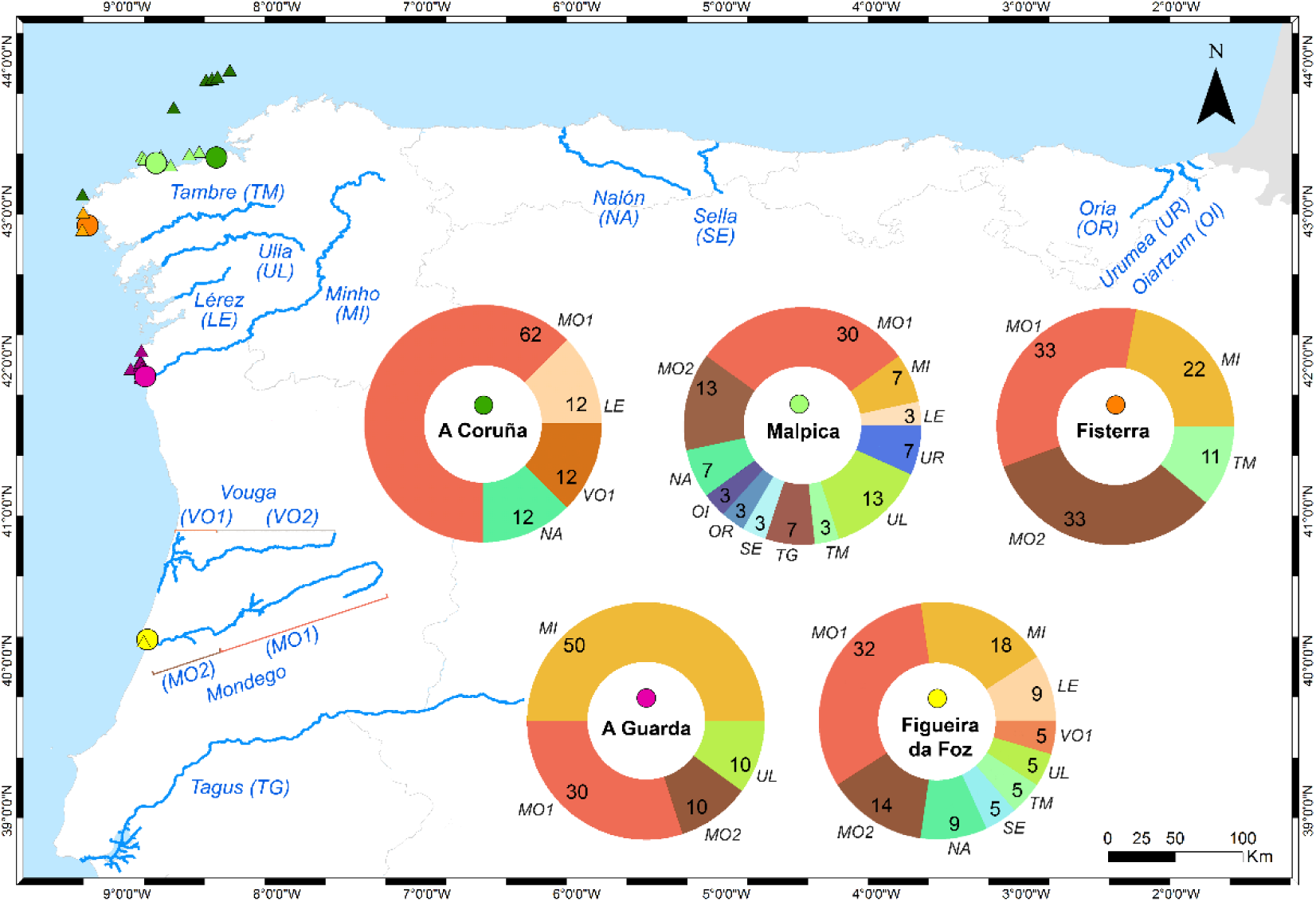
Natal-river assignments of adult bycatch Allis shad *Alosa alosa* (L. 1758) specimens at an 80 % assignment-probability threshold, displayed per fish market as donut charts. Donut slices are labelled with two-letter river abbreviations, and their angular widths correspond to the percentage of specimens assigned. River segments with assigned individuals appear in blue on the map; Vouga 2 is highlighted in grey as it’s without assignments in comparison with its adjacent stretch (Vouga 1). Triangles— colour-coded by fish market—mark the approximate marine capture locations. Due to the map’s scale and the catches’ proximity to shore, the symbols may visually overlap the coastline, though all points are correctly placed just offshore.

**Figure 8.**
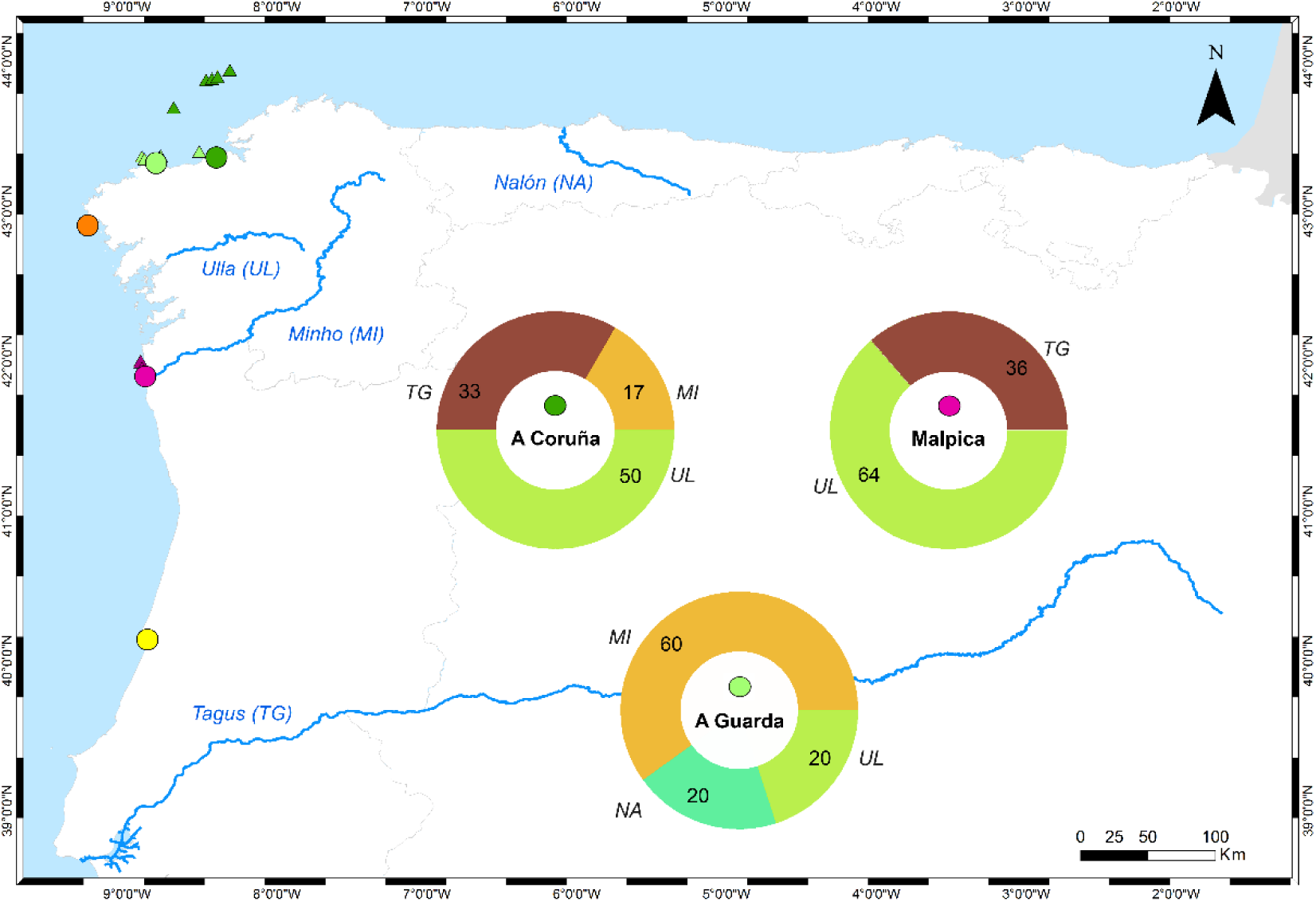
Natal-river assignments of adult bycatch Twaite shad *Alosa fallax* (Lacépede 1803) specimens at an 80 % assignment-probability threshold, displayed per fish market as donut charts. Donut slices are labelled with two-letter river abbreviations, and their angular widths correspond to the percentage of specimens assigned. River segments with assigned individuals appear in blue on the map. Triangles—colour-coded by fish market—mark the approximate marine capture locations. Due to the map’s scale and the catches’ proximity to shore, the symbols may visually overlap the coastline, though all points are correctly placed just offshore.

**Table 5.** Assignment of bycaught Allis shad *Alosa alosa* (L. 1758) specimens to their natal river (geographically ordered from north to south), shown by fish market. Values are reported as number of assigned individuals (*N*) and percentage (%).

| Rivers | Fish Markets |  |  |  |  |  |  |  |  |  |  |  |
| --- | --- | --- | --- | --- | --- | --- | --- | --- | --- | --- | --- | --- |
|  | A Coruña |  | Malpica |  | Fisterra |  | A Guarda |  | Figueira da Foz |  | Total |  |
|  | *N* | % | *N* | % | *N* | % | *N* | % | *N* | % | *N* | % |
| Bidasoa | 0 | 0% | 0 | 0% | 0 | 0% | 0 | 0% | 0 | 0% | 0 | 0% |
| Oiartzun | 0 | 0% | 1 | 3.33% | 0 | 0% | 0 | 0% | 0 | 0% | 1 | 1.27% |
| Urumea | 0 | 0% | 2 | 6.67% | 0 | 0% | 0 | 0% | 0 | 0% | 2 | 2.53% |
| Oria | 0 | 0% | 1 | 3.33% | 0 | 0% | 0 | 0% | 0 | 0% | 1 | 1.27% |
| Asón | 0 | 0% | 0 | 0% | 0 | 0% | 0 | 0% | 0 | 0% | 0 | 0% |
| Pas | 0 | 0% | 0 | 0% | 0 | 0% | 0 | 0% | 0 | 0% | 0 | 0% |
| Deva | 0 | 0% | 0 | 0% | 0 | 0% | 0 | 0% | 0 | 0% | 0 | 0% |
| Sella | 0 | 0% | 1 | 3.33% | 0 | 0% | 0 | 0% | 1 | 4.55% | 2 | 2.53% |
| Nalón | 1 | 12.5% | 2 | 6.67% | 0 | 0% | 0 | 0% | 2 | 9.09% | 5 | 6.33% |
| Eo | 0 | 0% | 0 | 0% | 0 | 0% | 0 | 0% | 0 | 0% | 0 | 0% |
| Tambre | 0 | 0% | 1 | 3.33% | 1 | 11.11% | 0 | 0% | 1 | 4.55% | 3 | 3.80% |
| Ulla | 0 | 0% | 4 | 13.33% | 0 | 0% | 1 | 10% | 1 | 4.55% | 6 | 7.60% |
| Lérez | 1 | 12.5% | 1 | 3.33% | 0 | 0% | 0 | 0% | 2 | 9.09% | 4 | 5.06% |
| Minho | 0 | 0% | 2 | 6.67% | 2 | 22.22% | 5 | 50% | 4 | 18.18% | 13 | 16.46% |
| Lima | 0 | 0% | 0 | 0% | 0 | 0% | 0 | 0% | 0 | 0% | 0 | 0% |
| Vouga1 | 1 | 12.5% | 0 | 0% | 0 | 0% | 0 | 0% | 1 | 4.55% | 2 | 2.53% |
| Vouga2 | 0 | 0% | 0 | 0% | 0 | 0% | 0 | 0% | 0 | 0% | 0 | 0% |
| Mondego1 | 5 | 62.5% | 9 | 30% | 3 | 33.33% | 3 | 30% | 7 | 31.82% | 27 | 34.18% |
| Mondego2 | 0 | 0% | 4 | 13.33% | 3 | 33.33% | 1 | 10% | 3 | 13.64% | 11 | 13.92% |
| Tagus | 0 | 0% | 2 | 6.67% | 0 | 0% | 0 | 0% | 0 | 0% | 2 | 2.53% |
| Total | 8 | 100% | 30 | 100% | 9 | 100% | 10 | 100% | 22 | 100% | 79 | 100% |

Abbreviations: *N*, number of individuals; %, percentage

**Table 6.** Assignment of bycaught Twaite shad *Alosa fallax* (Lacépède 1803) specimens to their natal river (geographically ordered from north to south), shown by fish market. Values are reported as number of assigned individuals (*N*) and percentage (%).

| Rivers | Fish markets |  |  |  |  |  |  |  |
| --- | --- | --- | --- | --- | --- | --- | --- | --- |
|  | A Coruña |  | Malpica |  | A Guarda |  | Total |  |
|  | *N* | % | *N* | % | *N* | % | *N* | % |
| Bidasoa | 0 | 0% | 0 | 0% | 0 | 0% | 0 | 0% |
| Oiartzun | 0 | 0% | 0 | 0% | 0 | 0% | 0 | 0% |
| Urumea | 0 | 0% | 0 | 0% | 0 | 0% | 0 | 0% |
| Oria | 0 | 0% | 0 | 0% | 0 | 0% | 0 | 0% |
| Asón | 0 | 0% | 0 | 0% | 0 | 0% | 0 | 0% |
| Pas | 0 | 0% | 0 | 0% | 0 | 0% | 0 | 0% |
| Deva | 0 | 0% | 0 | 0% | 0 | 0% | 0 | 0% |
| Sella | 0 | 0% | 0 | 0% | 0 | 0% | 0 | 0% |
| Nalón | 0 | 0% | 0 | 0% | 1 | 20% | 1 | 3.57% |
| Eo | 0 | 0% | 0 | 0% | 0 | 0% | 0 | 0% |
| Tambre | 0 | 0% | 0 | 0% | 0 | 0% | 0 | 0% |
| Ulla | 6 | 50% | 7 | 63.64% | 1 | 20% | 14 | 50% |
| Lérez | 0 | 0% | 0 | 0% | 0 | 0% | 0 | 0% |
| Minho | 2 | 16.67% | 0 | 0% | 3 | 60% | 5 | 17.86% |
| Lima | 0 | 0% | 0 | 0% | 0 | 0% | 0 | 0% |
| Vouga1 | 0 | 0% | 0 | 0% | 0 | 0% | 0 | 0% |
| Vouga 2 | 0 | 0% | 0 | 0% | 0 | 0% | 0 | 0% |
| Mondego1 | 0 | 0% | 0 | 0% | 0 | 0% | 0 | 0% |
| Modengo2 | 0 | 0% | 0 | 0% | 0 | 0% | 0 | 0% |
| Tagus | 4 | 33.33% | 4 | 36.36% | 0 | 0% | 8 | 28.57% |
| Total | 12 | 100% | 11 | 100% | 5 | 100% | 28 | 100% |
| Abbreviations: *N*, number of individuals; %, percentage |  |  |  |  |  |  |  |  |

**Table 7.**
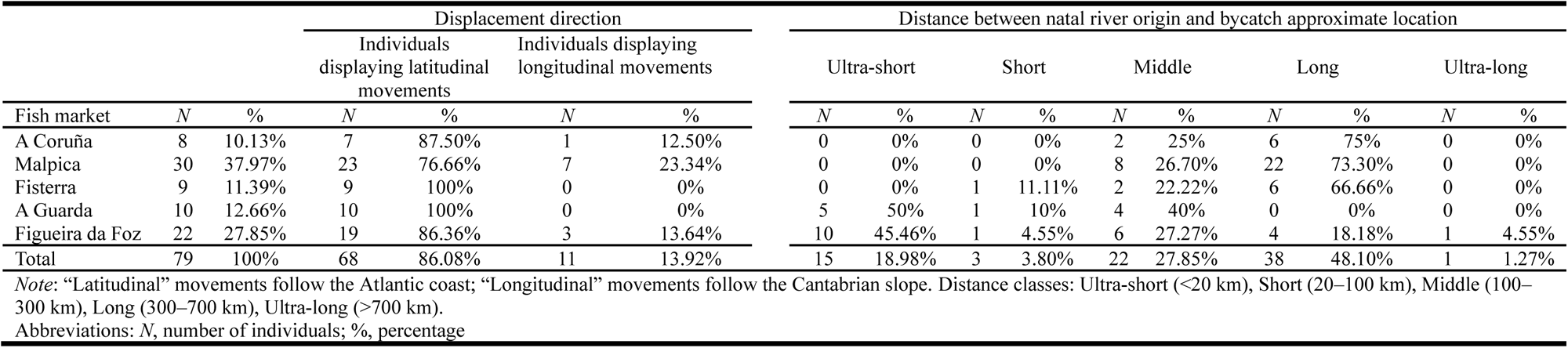
Direction and distance travelled by bycaught Allis shad *Alosa alosa* (L. 1758) from inferred natal rivers to the approximate marine bycatch-location centroid, grouped by fish market (north to south). Distances were measured as straight-line approximations (km) from the river mouth. Values are reported as number of assigned individuals (*N*) and percentage (%).

**Table 8.** Direction and distance travelled by bycaught Twaite shad *Alosa fallax* (Lacépède 1803) from inferred natal rivers to the approximate marine bycatch-location centroid, grouped by fish market (north to south). Distances were measured as straight-line approximations (km) from the river mouth. Values are reported as number of assigned individuals (*N*) and percentage (%).

|  |  |  | Displacement direction |  |  |  | Distance between natal river origin and bycatch approximate location |  |  |  |  |  |  |  |  |  |
| --- | --- | --- | --- | --- | --- | --- | --- | --- | --- | --- | --- | --- | --- | --- | --- | --- |
|  |  |  | Individuals displaying latitudinal movements |  | Individuals displaying longitudinal movements |  | Ultra-short |  | Short |  | Middle |  | Long |  | Ultra-long |  |
| Fish market | *N* | % | *N* | % | *N* | % | *N* | % | *N* | % | *N* | % | *N* | % | *N* | % |
| A Coruña | 12 | 42.86% | 12 | 100% | 0 | 0% | 0 | 0% | 0 | 0% | 8 | 66.7% | 4 | 33.3% | 0 | 0% |
| Malpica | 11 | 39.29% | 11 | 100% | 0 | 0% | 0 | 0% | 0 | 0% | 7 | 63.6% | 4 | 36.4% | 0 | 0% |
| A Guarda | 5 | 17.86% | 4 | 80% | 1 | 20% | 3 | 60% | 1 | 20% | 0 | 0% | 1 | 20% | 0 | 0% |
| Total | 28 | 100% | 27 | 96.43% | 1 | 3.57% | 3 | 10.71% | 1 | 3.57% | 15 | 53.57% | 9 | 32.14% | 0 | 0% |
| *Note:* “Latitudinal” movements follow the Atlantic coast; “Longitudinal” movements follow the Cantabrian slope. Distance classes: Ultra-short (<20 km), Short (20–100 km), Middle (100–300 km), Long (300–700 km), Ultra-long (>700 km).<br>Abbreviations: *N*, number of individuals; %, percentage |  |  |  |  |  |  |  |  |  |  |  |  |  |  |  |  |

## Discussion

To the best of our knowledge, this is the first study to explicitly link population mixing, marine dispersal capacities, and metapopulation functioning with bycatch vulnerability in European shads. Our results show that bycatch exposure is population-specific and structured by patterns of connectivity and dispersal, rather than random. Bycatch disproportionately impacts core populations, potentially altering metapopulation stability and source–sink dynamics. These findings underscore the importance of incorporating population structure and movement ecology into risk assessments and conservation planning for wide-ranging migratory species.

### Spatial patterns and species-specific patterns in natal origin of bycatch individuals

Our findings revealed spatial and species-specific differences in bycatch risk, as indicated by natal-origin diversity across marine areas and/or fishing markets. *Alosa alosa* exhibited the highest diversity of natal origins in Malpica and Figueira da Foz, while *A. fallax* peaked in A Coruña and A Guarda. Despite their geographical proximity, these critical bycatch zones showed clear species-specific segregation. Nevertheless, both species displayed maximal natal-origin diversity at opposite ends of the study area, suggesting a shared pattern of latitudinal coastal diffusion. This coastal diffusion pattern echoes findings in European shad populations along the French Atlantic coast (Nachón *et al*., 2020), as well as for other *Alosa* species, such as Alewife *Alosa pseudoharengus* (Wilson 1811) and Blueback herring *Alosa aestivalis* (Mitchill 1814) along the US Atlantic coast (Hasselman *et al*., 2016), and American shad *Alosa sapidissima* (Wilson 1811) along the coast of Maine, USA, and Nova Scotia, Canada (Walther and Thorrold, 2010). Such distributional patterns likely reflect a complex interplay of riverine proximity, trophic resource availability, and density-dependent processes (Sánchez-Hernández *et al*., 2017; Schlosser, 1998; Taverny and Elie, 2001; Thorrold *et al*., 2001), though the precise drivers warrant further investigation.

Clear interspecific differences in both natal origin diversity and dispersal patterns were evident. *Alosa alosa* exhibited greater natal-origin diversity and more extensive marine displacements than *A. fallax*. *A. alosa* more frequently undertook middle- and long-distance movements, including rare longitudinal displacements along the Cantabrian slope, whereas *A. fallax* was largely restricted to latitudinal movements over middle distances. This contrast aligns with previous findings indicating that *A. alosa* functions as a broader metapopulation, with lower philopatry and greater dispersal capacity than *A. fallax* (Jolly *et al*., 2012; Martin *et al*., 2015; Nachón *et al*., 2020; Randon *et al*., 2018; Rougemont *et al*., 2022; Sabatino *et al*., 2022; Taillebois *et al*., 2020), as well as a wider distribution, particularly along the Cantabrian coast (Aprahamian *et al*., 2015; Barber-O’Malley *et al*., 2022; Doadrio *et al*., 2011; Mota *et al*., 2016; Nachón *et al*., 2016, 2019a, b; Wilson and Veneranta, 2019). In general, the most frequent or dominant natal origins of *A. alosa* were more widely distributed across the study area than those of *A. fallax*. For *A. alosa*, Mondego and Minho origins were prevalent near the mouths of their respective rivers but were also detected throughout the entire coast—especially those from the Mondego River, which dominated all marine areas. By contrast, *A. fallax* displayed a more spatially structured pattern: Ulla River origins were most frequent in the northern marine areas near the Ulla estuary, while Minho River origins predominated in the southernmost area, near the Minho mouth. However, *A. fallax* individuals assigned to the Tagus River appeared in northern marine areas, suggesting that some *A. fallax* contingents may also undertake broader coastal dispersal. A similar duality in spatial behaviour appears to be present in both species, though more pronounced in *A. alosa*. These patterns reflect the dual resident–dispersive structure previously described by Nachón *et al*. (2020) for European shads along the Bay of Biscay (France), where both contingents coexisted within the same population. Certain assignments, however, demand cautious interpretation—notably *A. alosa* individuals attributed to the Ulla River, a system currently recognized as monospecific for *A. fallax* (Nachón *et al*., 2016). These cases may reflect microchemical overlap between populations rather than true anomalies in species distribution. Nonetheless, the broader patterns observed offer valuable insights into how bycatch pressure interacts with species-specific dispersal traits to structure population connectivity across marine space.

### Population mixing, metapopulation dynamics and management implications

The predominance of Mondego and Minho River origins in *Alosa alosa* bycatch, contrasted with Minho and Ulla River origins for *A. fallax*—consistent with these being recognized the most abundant populations for each species (Mota *et al*., 2015, 2016; Nachón *et al*., 2016; Stratoudakis *et al*., 2016)—suggests that these populations may play distinct roles within their respective metapopulation functioning. Interestingly, the Tagus River emerged as a relevant contributor to *A. fallax* bycatch—an unexpected finding given its distance from the fishing grounds and the assumed poor or uncertain status of its populations (Almeida *et al*., 2023; Costa *et al*., 2001)—underscoring the importance of further investigating marine population mixing and metapopulation dynamics. Within a metapopulation dynamics—particularly relevant for *A. alosa* (Randon *et al*., 2018) and likely operating at a more regional scale for *A. fallax* (Jolly *et al*., 2012; Martin *et al*., 2015; Sabatino *et al*., 2022)—these more abundant populations likely function as source populations, exporting individuals to sink populations through marine dispersal. Their higher abundance may drive a diffusion effect, whereby a greater absolute number of emigrants leads to broader spatial dispersion. This diffusion effect has been previously identified in the French coast of the Bay of Biscay, where the dominant European shad continental populations in the 1980s exhibited substantial marine dispersion due to their higher abundance (Nachón *et al*., 2020). Consequently, these populations are more frequently encountered by fisheries, increasing their susceptibility to bycatch. Although these populations are supposed to be relatively abundant, the disproportionate bycatch pressure raises concerns about their long-term viability. Smaller or less abundant populations, with their limited numbers, may be more vulnerable to local extirpation due to bycatch. However, if the expansion of more successful natal origins is contributing to the spread of individuals into smaller populations, this could result in a natural mixing process, where larger populations colonize the rivers of less abundant stocks.

However, it is important to note that bycatch pressure is not always proportional to the relative abundance of populations. For example, Hasselman *et al*. (2016) found that bycatch mortality in other *Alosa* species, such as *A. pseudoharengus* and *A. aestivalis*, was disproportionately assigned to more depleted populations. This suggests that more abundant populations are not necessarily the most affected by bycatch. Rather, it is those populations in a more vulnerable state that may suffer the most significant impacts, regardless of their relative abundance. The relationship between bycatch and population depletion remains complex, and it is unclear whether these populations were already in a depleted state before the study or whether bycatch is contributing to their decline. These uncertainties further highlight the need for careful consideration of the status and dynamics of populations when evaluating bycatch impacts. Studies on species with complex population structures, such as Atlantic cod *Gadus morhua* L. 1758 in the Baltic Sea, show that mixing between different populations complicates stock assessments and fisheries management (Hüssy *et al*., 2016). Treating distinct cod stocks as a single unit can mask their different dynamics, highlighting the need for tailored management strategies, particularly for species like European shads with metapopulation structures. Likewise, Puncher *et al*. (2018) discussed similar challenges in managing migratory species such as Atlantic bluefin tuna *Thunnus thynnus* (L. 1758), which exhibit population differentiation across wide geographical areas. These examples underscore the importance of considering spatial dynamics and population structure when developing effective fisheries management strategies, especially for species like shad, where bycatch can significantly impact long-term population health.

These species-specific differences in bycatch composition have important implications for conservation and management. Notably, the more constrained spatial distribution of *A. fallax*, with stronger philopatry, suggests that this species may be more vulnerable to localized bycatch pressure, particularly in regions close to its natal rivers. Conversely, *A. alosa*’s broader dispersal and higher natal-origin diversity across the fishing zones indicate a more widespread vulnerability to bycatch, potentially impacting populations at a regional scale. This contrast highlights the complexity of bycatch risk management: species with greater dispersal may be affected by fishing pressures across larger areas, while more philopatric species may face disproportionate impacts in specific zones. Furthermore, *A. alosa* is less polymorphic, phenotypically plastic, and more vulnerable to anthropogenic pressures throughout its life cycle than *A. fallax* (Baglinière, 2000), contributing to its poorer conservation status in the study countries (Mota *et al*., 2016; Nachón *et al*., 2016, 2019a, b; Stratoudakis *et al*., 2016). Bycatch impacts are unevenly distributed, with the most affected populations being those of *A. alosa* from the Mondego and Minho rivers. These populations are critical for both ecological and socioeconomic reasons, as they are among the last to support artisanal inland fisheries (Almeida *et al*., 2023; Azeiteiro *et al*., 2021; Braga *et al*., 2022; Stratoudakis *et al*., 2023). To remain a viable species for commercial exploitation in the region, it is essential to implement management measures that consider both the spatial distribution of bycatch risk and the vulnerability of specific populations. The restoration efforts in the Mondego River, which have improved connectivity and facilitated access to upstream habitats (Belo *et al*., 2021), demonstrate the potential for recovery. However, incidental capture threatens to reverse these advances by removing individuals born in recolonized habitats, as observed in our study. Consequently, integrated management strategies are essential to ensure the long-term recovery and sustainability of shad populations.

### Methodological framework: considerations and future directions

Our findings should be interpreted within the strategic framework of this study’s design, which prioritized spatial and biological relevance during a critical life-history phase. By aligning sampling with the onset of spawning migration, when adult shads aggregate near major river mouths (La Mesa *et al*., 2015; Nachón *et al*., 2016; Taverny and Elie, 2001; Trancart *et al*., 2014), we aimed to capture population connectivity during this ecologically pivotal period. The ∼400 km coastal coverage focused on key bycatch hotspots and fish markets—areas where fishing activity directly overlaps with essential European shad marine habitats (Nachón *et al*., 2022a, b, c, d)—particularly near the Ulla, Minho, and Mondego rivers, which sustain the region’s most stable populations of both species (Belo *et al*., 2021; Mota *et al*., 2015; Nachón *et al*., 2016; Stratoudakis *et al*., 2016, 2023).

For geochemical assignments, we leveraged the diagnostic power of ^87^Sr/^86^Sr ratios as ideal spatial markers, given their ability to reflect distinct bedrock geologies while remaining stable over ecological timescales and precisely reflecting stream water values without trophic fractionation (Kennedy *et al*., 2000; Loewen *et al*., 2015; Martin *et al*., 2013a; Zimmerman *et al*., 2013). Complementary Sr/Ca and Ba/Ca ratios enhanced discrimination precision, building on established methodologies (Fontaine *et al*., 2025; Martin *et al*., 2013b, 2015; Nachón *et al*., 2020; Randon *et al*., 2018). Our fluvial baseline was specifically designed to capture the regular dispersal ecology of both species (Jolly *et al*., 2012; Martin *et al*., 2015; Nachón *et al*., 2020; Randon *et al*., 2018; Sabatino *et al*., 2022), ensuring reliable discrimination of natal origins. At regional scales, the marked geological contrast between the granitic Atlantic coast (^87^Sr/^86^Sr >0.715) and calcareous/sedimentary Cantabrian coast (^87^Sr/^86^Sr <0.710) provided robust spatial resolution. At finer scales, while natal river differentiation can be more challenging due to overlapping of ^87^Sr:^86^Sr chemical signatures, the combined use of isotopic (^87^Sr/^86^Sr) and elemental tracers (Sr/Ca and Ba/Ca ratios) allowed for a reliable assignment, particularly for core natal systems like the Ulla, Minho, and Mondego rivers.

Conservative probability thresholds and QDA-based classification—selected for its balance between computational efficiency and performance—yielded high assignment accuracy (94.76–96.83%). This high precision was maintained even considering potential chemical signature overlap in some regions, one of the primary uncertainties in natal river assignments (Fontaine *et al*., 2025). Collectively, this approach provides a robust yet nuanced framework for evaluating broad-scale bycatch impacts across distinct natal origins during critical life stages. However, given the inherent variability in geochemical signatures across river basins (Bareille *et al*., 2024; Martin *et al*., 2013a) and the spatial dispersal of individuals throughout their life cycle (Jolly *et al*., 2012; Martin *et al*., 2015; Nachón *et al*., 2020; Randon *et al*., 2018; Sabatino *et al*., 2022), continued refinement of these methods is necessary to further improve resolution and assignment confidence.

Our findings offer a concise snapshot of a complex and dynamic fisheries management issue. Future research should focus on developing quantitative models that link bycatch patterns to riverine population distribution, structure, and abundance. The use of non-invasive tools such as environmental DNA (eDNA), particularly valuable for detecting rare or threatened migratory species (Antognazza *et al*., 2019, 2021; Bhendarkar *et al*., 2025), could enhance our ability to establish such links between freshwater distribution and/or abundance and marine dispersal. Expanding the spatial and temporal scope of studies and investigating differences across size and age classes as well as fishing gear types, could help assess how bycatch impacts vary across regions, seasons, and life-history stages, thereby refining management strategies. Additionally, given the complex hybridization dynamics between these sister species (Taillebois *et al*., 2020; Rougemont *et al*., 2022), future studies should evaluate the impact of bycatch on hybrid individuals, which have been observed alongside *A. alosa* and *A. fallax* along the Galician coast (Nachón *et al*., 2022a, b, c, d). Understanding the natal origins and ecological role of hybrids will be crucial for assessing their conservation status and adaptive potential. Ultimately, integrating these insights into spatially explicit conservation strategies will be key to mitigating bycatch effects. This includes implementing targeted management measures that account for population-specific dispersal patterns and bycatch hotspots.

## Conclusion

This study highlights how bycatch in European shads is shaped by natal origin and dispersal behaviour, disproportionately affecting individuals from core, productive populations and areas of high natal diversity, particularly in estuarine and coastal zones under intense fishing pressure. These patterns involve exchanges between riverine and marine populations across different countries, transcending political boundaries. Given the widespread nature of bycatch across the species’ distribution range (ICES, 2014; King and Roche, 2008; La Mesa *et al*., 2015; Nachón *et al*., 2016; Sabatié, 1993; Trancart *et al*., 2014), this study provides a foundation for expanding similar approaches to other regions and emphasizes the need for transnational, ecosystem-based management like DiadES (2024) and DiadSea Interreg Atlantic Area project initiatives. Effective conservation must integrate freshwater and marine components—often overlooked in current frameworks (Verhelst *et al*., 2021)—and prioritize critical habitats such as river mouths and feeding grounds. Habitat-based functionality models, linking population functionality with habitat suitability (Dambrine *et al*., 2023), and mechanistic species distribution models such as GR3D, which account for straying, reproductive strategies, and oceanic distribution (Poulet *et al*., 2023), offer promising avenues to improve bycatch mitigation through natal-origin-informed restrictions. Initiatives like voluntary avoidance programs emphasize the importance of fisher participation in bycatch reduction (Bethoney *et al*., 2017), with cooperation among fishermen, ecologists, and authorities critical for sustainable fisheries, especially in areas with limited fisher organization (Stratoudakis *et al*., 2020).

## Acknowledgements

This work was supported by the project DiadES—Assessing and enhancing ecosystem services provided by diadromous fish in a climate change context (EAPA_18/2018), funded by the INTERREG Atlantic Area programme. Additional funding was provided by the 1MARDEALOSAS project, supported by the Biodiversity Foundation of the Ministry for Ecological Transition and the Demographic Challenge, through the Pleamar Programme, co-financed by the European Maritime and Fisheries Fund (EMFF). We are grateful to the fish markets and fishermen’s associations and fishermen that collaborated in this study: A Coruña, Malpica, Fisterra, A Guarda, and Figueira da Foz. Special thanks to their representatives, who facilitated the acquisition of specimens, mediated with fishermen, and addressed logistical issues related to fish storage: Nacho (A Coruña), Francisco (Malpica), Carlos (Fisterra), Manuel (A Guarda) and Júlio (Figueira da Foz). We also thank Patrick Lambert and Philippe Jatteau at INRAE (Cestas, Bordeaux) for generously providing A. fallax juvenile otoliths from systematic sampling in the Garonne and Dordogne basins, housed in the INRAE otolith collection.

## Contributions

**D. J. N.:** Conceptualization, methodology, validation, formal analysis, investigation, data curation, writing–original draft, writing–review and editing, visualization, project administration, supervision; **A. P.:** Methodology, validation, formal analysis, data curation, visualization, writing–review and editing; **F. D.:** Conceptualization, methodology, investigation, writing–review and editing, supervision, project administration; **R. V-L.:** Investigation, visualization, writing–review and editing; **R. C.:** Validation, writing–review and editing; **A. F. B.:** Investigation, data curation, writing– review and editing; **C. S. M.:** Investigation, writing–review and editing; **B. R. Q.:** Investigation, writing–review and editing; **P. R. A.:** Writing–review and editing, funding acquisition, project administration; **C. A.:** Investigation, funding acquisition; **G. B.:** Validation, investigation, writing–review and editing; **C. P.:** Investigation, validation, resources; **F. Cl.:** Investigation, validation, resources; **P. L.:** Writing–review and editing, funding acquisition, project administration; **G. L.:** Writing–review and editing, funding acquisition, project administration, supervision; **F. C.:** Conceptualization, funding acquisition, project administration, supervision. All authors have read and approved the final version of the manuscript.

## Funding information

This work was supported by the project *DiadES—Assessing and enhancing ecosystem services provided by diadromous fish in a climate change context* (EAPA_18/2018), funded by the INTERREG Atlantic Area program. Additional funding was provided by the project *1MARDEALOSAS—Evaluación de las “capturas incidentales” de* Alosa alosa *y* Alosa fallax *por la flota costera de Galicia: análisis del problema, sensibilización y proposición de medidas de gestión y protección*, supported by the Biodiversity Foundation of the Ministry for Ecological Transition and the Demographic Challenge, through the Pleamar Programme, co-financed by the European Maritime and Fisheries Fund (EMFF). This work was also supported by Portuguese National Funds through FCT — Foundation for Science and Technology through MARE’s base funding (UIDB/04292/2020, https://doi.org/10.54499/UIDB/04292/2020) and MARE’s strategic program (https://doi.org/10.54499/UIDP/04292/2020), through project LA/P/0069/2020 granted to the Associate Laboratory ARNET (https://doi.org/10.54499/LA/P/0069/2020) and through the FCT doctoral grant attributed to Ana F. Belo (SFRH/BD/123434/2016).

## Notes

https://doi.org/10.5281/zenodo.15462157

### Competing Interest Statement

The authors have declared no competing interest.

## References

Alexandrino, P., Faria, R., Linhares, D., Castro, F., Le Corre, M., Sabatié, R., Baglinière, J. L., & Weiss, S. (2006). Interspecific differentiation and intraspecific substructure in two closely related clupeids with extensive hybridization, *Alosa alosa* and *Alosa fallax*. Journal of Fish Biology, 69, 242–259. 10.1111/j.1095-8649.2006.01289.x

Almeida, P. R., Mateus, C. S., Alexandre, C. M., Pedro, S., Boavida-Portugal, J., Belo, A. F., Pereira, E., Silva, S., Oliveira, I., & Quintella, B. R. (2023). The decline of the ecosystem services generated by anadromous fish in the Iberian Peninsula. Hydrobiologia, 850(12), 2927–2961. 10.1007/s10750-023-05179-6

Anonymous (2025). Data and codes of the manuscript “Population-specific bycatch risks in two vulnerable anadromous clupeids: insights from otolith microchemistry.” Zenodo Digital Repository, (V2.0). 10.5281/zenodo.15462157

Antognazza, C. M., Britton, J. R., De Santis, V., Kolia, K., Turunen, O. A., Davies, P., Allen, L., Hardouin, E. A., Crundwell, C., & Andreou, D. (2021). Environmental DNA reveals the temporal and spatial extent of spawning migrations of European shad in a highly fragmented river basin. Aquatic Conservation: Marine and Freshwater Ecosystems, 31(8), 2029–2040. 10.1002/aqc.3601

Antognazza, C. M., Britton, J. R., Potter, C., Franklin, E., Hardouin, E. A., Gutmann Roberts, C., Aprahamian, M., & Andreou, D. (2019). Environmental DNA as a non-invasive sampling tool to detect the spawning distribution of European anadromous shads (*Alosa* spp.). Aquatic Conservation: Marine and Freshwater Ecosystems, 29(1), 148–152. 10.1002/aqc.3010

Aprahamian, M., Alexandrino, P., Antunes, C., Cobo, F., King, J., Lambert, P., Martin, J., Mota, M., Nachón, D. J., & Silva, S. (2015). Shads state of the art. In P. R. Almeida & E. Rochard (Eds.), ICES Report of the Workshop on Lampreys and Shads (WKLS), 27–29 November 2014, Lisbon, Portugal, ICES CM 2014/SSGEF:13 (pp. 46–105). International Council for the Exploration of the Sea, Copenhagen. Available at: https://ices-library.figshare.com/articles/report/Report_of_the_Workshop_on_Lampreys_and_Shads_WKLS_/18613613?file=33391082 (last accessed 15 April 2025).

Aprahamian, M. W., Aprahamian, C. D., Baglinière, J. L., Sabatié, R., & Alexandrino, P. (2003). Alosa alosa and Alosa fallax spp.: Literature review and bibliography. (R&D Technical Report W1-014/TR). Environment Agency, Bristol. Available at: https://assets.publishing.service.gov.uk/media/5a7c543840f0b62dffde1617/sw1-014-tr-e-e.pdf (last accessed 15 April 2025).

Aprahamian, M. W., Aprahamian, C. D., & Knights, A. M. (2010). Climate change and the green energy paradox: the consequences for twaite shad *Alosa fallax* from the River Severn, UK. Journal of Fish Biology, 77(8), 1912–1930. 10.1111/j.1095-8649.2010.02775.x

Ashley, M., Murillas, A., Muench, A., Marta-Pedroso, C., Rodwell, L., Rees, S., Rendle, E., Bašić, T., Copp, G. H., Díaz, E., Nachón, D. J., Lambert, P., & Lassalle, G. (2023). An evidence base of ecosystems services provided by diadromous fish in the European Atlantic Area. Ecosystem Services, 64, 101559. 10.1016/j.ecoser.2023.101559

Azeiteiro, U. M., Pereira, M. J., Soares, A. M., Braga, H. O., Morgado, F., Sousa, M. C., Días, J. M., & Antunes, C. (2021). Dynamics of two anadromous species in a dam intersected river: analysis of two 100-year datasets. Fishes, 6(2), 21. 10.3390/fishes6020021

Baglinière, J. L. (2000). Le genre *Alosa* sp. In J. L. Baglinière & P. Elie (Eds.), Les aloses (Alosa alosa et Alosa fallax spp.). Écobiologie et variabilité des populations (pp. 3–30). Paris: CEMAGREF-INRA Editions.

Baglinière, J. L., Sabatié, M. R., Rochard, E., Alexandrino, P., & Aprahamian, M. W. (2003). The allis shad *Alosa alosa*: Biology, ecology, range, and status of populations. American Fisheries Society Symposium, 35, 85–102.

Barber-O’Malley, B., Lassalle, G., Lambert, P., & Quinton, E. (2022). Dataset on European diadromous species distributions from 1750 to present time in Europe, North Africa and the Middle East. Data in Brief, 40, 107821. 10.1016/j.dib.2022.107821

Bareille, G., Vignon, M., Chappaz, A., Fontaine, A., Tabouret, H., Morat, F., Martin, J., Aymes, J.C., Daverat, F., Pécheyran, C., & Donard, O. (2024). Freshwater fish otoliths record signals from both water and physiological processes: new insights from Sr/Ca and Ba/Ca ratios. Canadian Journal of Fisheries and Aquatic Sciences, 81, 223–240. 10.1139/cjfas-2022-0030

Barreiro, M., Abad, E., Antelo, L. T., Fernández, J. C., Pereira, C., Ovalle, J. C., Pérez Martín, R. I., & Valeiras, J. (2025). Development of smart electronic observation onboard technologies for more sustainable fisheries management. Frontiers in Marine Science, 12, 1545718. 10.3389/fmars.2025.1545718

Belo, A. F., Cardoso, G., Pereira, E., Quintella, B. R., Mateus, C. S., Alexandre, C. M., Batista, C., Telhado, A., Quadrado, M. F., & Almeida, P. R. (2021). Fish pass use by shads (*Alosa alosa* L. and *Alosa fallax* [Lacépède, 1803]): Implications for monitoring and management. Ecohydrology, 14(5), e2292. 10.1002/eco.2292

Bethoney, N. D., Schondelmeier, B. P., Kneebone, J., & Hoffman, W. S. (2017). Bridges to best management: effects of a voluntary bycatch avoidance program in a mid-water trawl fishery. Marine Policy, 83, 172–178. 10.1016/j.marpol.2017.06.003

Bhendarkar, M., Claver, C., Mendibil, I., Fraija-Fernández, N., Nachón, D. J., Davison, P. I., Bašić, T., O’Leary, C., Roche, W. K., Azpiroz, I., Acolas, M-L., Lekuona, A., Ardaiz, J., Díaz, E., Lambert, P., Lassalle, G., & Rodríguez-Ezpeleta, N. (2025). Lessons learned from applying eDNA surveying to diadromous fish detection across the north-east Atlantic region. bioRxiv, 2025–01. 10.1101/2025.01.31.635873

Braga, H. O., Bender, M. G., Oliveira, H. M., Pereira, M. J., & Azeiteiro, U. M. (2022). Fishers’ knowledge on historical changes and conservation of Allis shad-*Alosa alosa* (Linnaeus, 1758) in Minho River, Iberian Peninsula. Regional Studies in Marine Science, 49, 102094. 10.1016/j.rsma.2021.102094

Campana, S. E. (1999). Chemistry and composition of fish otoliths: pathways, mecanisms and applications. Marine Ecology Progress Series, 188, 262–297. 10.3354/meps188263

Campana, S. E., & Thorrold, S. R. (2001). Otoliths, increments, and elements: keys to a comprehensive understanding of fish populations? Canadian Journal of Fisheries and Aquatic Sciences, 58(1), 30–38. doi:10.1139/f00-177

Chuenpagdee, R., Morgan, L. E., Maxwell, S. M., Norse, E. A., & Pauly, D. (2003). Shifting gears: assessing collateral impacts of fishing methods in US waters. Frontiers in Ecology and the Environment, 1(10), 517–524. 10.1890/1540-9295(2003)001[0517:SGACIO]2.0.CO;2

Cohen, J. (1988). Statistical power analysis for the behavioral sciences (2nd ed.). New York, NY: Lawrence Erlbaum Associates.

Costa, M. J., Almeida, P. R., Domingos, I. M., Costa, J. L., Correia, M. J., Chaves, M. L., & Teixeira, C. M. (2001). Present status of the main shads’ populations in Portugal. Bulletin Français de la Pêche et de la Pisciculture, (362-363), 1109–1116. 10.1051/kmae:2001040

Dambrine, C., Lambert, P., Elliott, S., Boavida-Portugal, J., Mateus, C. S., O’Leary, C., Pauwels, I., Poole, R., Roche, W., Van den Bergh, E., Vanoverbeke, J., & Lassalle, G. (2023). Connecting population functionality with distribution model predictions to support freshwater and marine management of diadromous fish species. Biological Conservation, 287, 110324. 10.1016/j.biocon.2023.110324

Daverat, F., & Martin, J. (2016). Microchemical and schlerochronological analyses used to infer fish migration. In P. Morais & F. Daverat (Eds.), An Introduction to Fish Migration (pp. 149–168). Boca Raton: CRC Press. 10.1201/b21321

Daverat, F., Martin, J., Fablet, R., & Pécheyran, C. (2011). Colonisation tactics of three temperate catadromous species, eel *Anguilla anguilla*, mullet *Liza ramada* and flounder *Plathychtys flesus*, revealed by Bayesian multielemental otolith microchemistry approach. Ecology of Freshwater Fish, 20, 42–51. 10.1111/j.1600-0633.2010.00454.x

Davies, P., Britton, R. J., Nunn, A. D., Dodd, J. R., Crundwell, C., Velterop, R., Ó’Maoiléidigh, N., O’Neill; R., Sheehan, V., Stamp, T., & Bolland, J. D. (2020). Novel insights into the marine phase and river fidelity of anadromous twaite shad *Alosa fallax* in the UK and Ireland. Aquatic Conservation: Marine and Freshwater Ecosystems, 30(7), 1291–1298. 10.1002/aqc.3343

Davies, R. W., Cripps, S. J., Nickson, A., & Porter, G. (2009). Defining and estimating global marine fisheries bycatch. Marine Policy, 33(4), 661–672. 10.1016/j.marpol.2009.01.003

DiadES (2024). A Transnational and Non-Sectoral Approach to Diadromous Fish Species Management. Available at: https://diades.eu/wp-content/uploads/2024/03/DiadES_policy_brief.pdf (last accessed 16 April 2025)

DiadSea Interreg Atlantic Area project. DiadSea project. Available at: https://www.atlanticarea.eu/discover-our-projects/approved-projects/diadsea (last accessed 15 April 2025).

Doadrio, I., Perea, S., Garzón-Heydt, P., & González, J. L. (2011). Ictiofauna continental española. Bases para su seguimiento. Madrid: DG Medio Natural y Política Forestal, MARM.

Elie, P., Taverny, C., Mennesson-Boisneau, C., & Sabatié, M. R. (2000). L’exploitation halieutique. In J. L. Baglinière & P. Elie (Eds.), Les aloses (Alosa alosa et Alosa fallax spp.). Écobiologie et variabilité des populations (pp. 199–226). Paris: CEMAGREF-INRA Editions.

Friedman, J. H. (1989). Regularized discriminant analysis. Journal of the American statistical association, 84(405), 165–175. 10.1080/01621459.1989.10478752

Fontaine, A., Vignon, M., Tabouret, H., Holub, A., Barranco, G., Bosc, S., Caux, I., Nachón, D. J., Elso, J., Caballero, P., Pécheyran, C., & Bareille, G. (2025). Inter-annual dispersal stability within the Atlantic salmon metapopulation from the Bay of Biscay. Fisheries Research, 284, 107323. 10.1016/j.fishres.2025.107323

Gray, C. A., & Kennelly, S. J. (2018). Bycatches of endangered, threatened and protected species in marine fisheries. Reviews in Fish Biology and Fisheries, 28(3), 521–541. 10.1007/s11160-018-9520-7

Hall, C. J., Jordaan, A., & Frisk, M. G. (2012). Centuries of anadromous forage fish loss: consequences for ecosystem connectivity and productivity. BioScience, 62(8), 723–731. 10.1525/bio.2012.62.8.5

Hasselman, D. J., Anderson, E. C., Argo, E. E., Bethoney, N. D., Gephard, S. R., Post, D. M., Schondelmeier, B. P., Schultz, T. F., Willis, T. V., & Palkovacs, E. P. (2016). Genetic stock composition of marine bycatch reveals disproportional impacts on depleted river herring genetic stocks. Canadian Journal of Fisheries and Aquatic Sciences, 73(6), 951–963. 10.1139/cjfas-2015-0402

Hastie, T., Tibshirani, R., & Friedman, J. (2009). The Elements of Statistical Learning: Data Mining, Inference, and Prediction (2nd ed.). New York, NY: Springer. 10.1007/978-0-387-84858-7

Hazen, E. L., Scales, K. L., Maxwell, S. M., Briscoe, D. K., Welch, H., Bograd, S. J., Bailey, H., Benson, S. R., Eguchi, T., Suzy, H. D., Costa, D. P., Crowder, L. B., & Lewison, R. L. (2018). A dynamic ocean management tool to reduce bycatch and support sustainable fisheries. Science advances, 4(5), eaar3001. 10.1126/sciadv.aar3001

Hüssy, K., Hinrichsen, H. H., Eero, M., Mosegaard, H., Hemmer-Hansen, J., Lehmann, A., & Lundgaard, L. S. (2016). Spatio-temporal trends in stock mixing of eastern and western Baltic cod in the Arkona Basin and the implications for recruitment. ICES Journal of Marine Science, 73(2), 293–303. 10.1093/icesjms/fsv227

ICES (2014). Report of the Working Group on Bycatch of Protected Species (WGBYC). Report ICES CM 2014/ACOM:28, Copenhagen. Available at: https://ices-library.figshare.com/articles/report/Report_of_the_Working_Group_on_Bycatch_of_Protected_Species_WGBYC_/19282769 (last accessed 15 April 2025).

ICES (2015). Report of the Workshop on Lampreys and Shads (WKLS), ICES CM 2014/SSGEF:13, Lisbon. Available at: https://ices-library.figshare.com/articles/report/Report_of_the_Workshop_on_Lampreys_and_Shads_WKLS_/18613613 (last accessed 15 April 2025).

ICES (2022). ICES Roadmap for bycatch advice on protected, endangered and threatened species (2022). ICES Technical Guidelines. Report. Available at: https://ices-library.figshare.com/articles/report/ICES_Roadmap_for_bycatch_advice_on_protected_endangered_and_threatened_species_2022_/19657167/1?file=35854196 (last accessed 15 April 2025).

Jolly, M. T., Aprahamian, M. W., Hawkins, S. J., Henderson, P. A., Hillman, R., O’Maoiléidigh, N., Maitland, P. S., Piper, R., & Genner, M. J. (2012). Population genetic structure of protected allis shad (*Alosa alosa*) and twaite shad (*Alosa fallax*). Marine Biology, 159, 675–687. 10.1007/s00227-011-1845-x

Kalish, J. M. (1989). Otolith microchemistry: validation of the effects of physiology, age and environment on otolith composition. Journal of Experimental Marine Biology and Ecology, 132, 151–178. 10.1016/0022-0981(89)90126-3

Kennedy, B. P., Blum, J. D., Folt, C. L., & Nislow, K. H. (2000). Using natural strontium isotopic signatures as fish markers: methodology and application. Canadian Journal of Fisheries and Aquatic Sciences, 57(11), 2280–2292. 10.1139/f00-206

King, J. J., & Roche, W. K. (2008). Aspects of anadromous Allis shad (*Alosa alosa* Linnaeus) and Twaite shad (*Alosa fallax* Lacépède) biology in four Irish Special Areas of Conservation (SACs): Status, spawning indications and implications for conservation designation. In S. Dufour, E. Prévost, E. Rochard, & P. Williot (Eds.), Fish and diadromy in Europe: Ecology, management, conservation (pp. 145–154). Springer Netherlands. 10.1007/978-1-4020-8548-2_15

La Mesa, G., Annunziatellis, A., Filidei Jr, E., & Fortuna, C. M. (2015). Modeling environmental, temporal and spatial effects on twaite shad (*Alosa fallax*) by-catches in the central Mediterranean Sea. Fisheries Oceanography, 24(2), 107–117. 10.1111/fog.12093

Lance, G. N., & Williams, W. T. (1966). Computer programs for hierarchical polythetic classification (’similarity analysis’). The Computer Journal, 9(1), 60–64. 10.1093/comjnl/9.1.60

Lance, G. N., & Williams, W. T. (1967). A general theory of classificatory sorting strategies: 1. Hierarchical systems. The Computer Journal, 9(4), 373–380. 10.1093/comjnl/9.4.373

Lassalle, G., Trancart, T., Lambert, P., & Rochard, E. (2008). Latitudinal variations in age and size at maturity among allis shad *Alosa alosa* populations. Journal of Fish Biology, 73(7), 1799–1809. 10.1111/j.1095-8649.2008.02036.x

Limburg, K. E., & Waldman, J. R. (2009). Dramatic declines in North Atlantic diadromous fishes. BioScience, 59(11), 955–965. 10.1525/bio.2009.59.11.7

Loewen, T. N., Reist, J. D., Yang, P., Koleszar, A., Babaluk, J. A., Mochnacz, N., & Halden, N. M. (2015). Discrimination of northern form Dolly Varden Char (*Salvelinus malma malma*) stocks of the North Slope, Yukon and Northwest Territories, Canada via otolith trace elements and ^87^Sr/^86^Sr isotopes. Fisheries Research, 170, 116–124. 10.1016/j.fishres.2015.05.025

Maitland, P. S., & Lyle, A. A. (2005). Ecology of Allis shad *Alosa alosa* and Twaite shad *Alosa fallax* in the Solway Firth, Scotland. Hydrobiologia, 534, 205–221. 10.1007/s10750-004-1571-1

Martin, J., Bareille, G., Bérail, S., Pécheyran, C., Daverat, F., Bru, N., Tabouret, H., & Donard, O. (2013a). Spatial and temporal variations in otolith chemistry and relationships with water chemistry: a useful tool to distinguish Atlantic salmon *Salmo salar* parr from different natal streams. Journal of Fish Biology, 82(5), 1556–1581. 10.1111/jfb.12089

Martin, J., Bareille, G., Berail, S., Pécheyran, C., Gueraud, F., Lange, F., Daverat, F., Bru, N., Beall, E., Barracou, D., & Donard, O. (2013b). Persistence of a southern Atlantic salmon population: diversity of natal origins from otolith elemental and Sr isotopic signatures. Canadian Journal of Fisheries and Aquatic Sciences, 70, 182–197. 10.1139/cjfas-2012-0284

Martin, J., Rougemont, Q., Drouineau, H., Launey, S., Jatteau, P., Bareille, G., Berail, S., Pécheyran, C., Feunteun, E., Roques, S., Clavé, D., Nachón, D. J., Antunes, C., Mota, M., Réveillac, E., & Daverat, F. (2015). Dispersal capacities of anadromous Allis shad population inferred from a coupled genetic and otolith approach. Canadian Journal of Fisheries and Aquatic Sciences, 72(7), 991–1003. 10.1139/cjfas-2014-0510

McDowall, R. M. (1988). Diadromy in fishes: Migrations between freshwater and marine environments. Londres, Uk: Croom Helm.

McDowall, R. M. (2001). Anadromy and homing: two life-history traits with adaptive synergies in salmonid fishes? Fish and Fisheries, 2(1), 78–85. 10.1046/j.1467-2979.2001.00036.x

Mota, M., Bio, A., Bao, M., Pascual, S., Rochard, E., & Antunes, C. (2015). New insights into biology and ecology of the Minho River Allis shad (*Alosa alosa* L.): contribution to the conservation of one of the last European shad populations. Reviews in Fish Biology and Fisheries, 25, 395–412. 10.1007/s11160-015-9383-0

Mota, M., Rochard, E., & Antunes, C. (2016). Status of the diadromous fish of the Iberian Peninsula: Past, present and trends. Limnetica, 35(1), 1–18. 10.23818/limn.35.01

Nachón, D. J. (2017). Dinámica poblacional y microquímica de los otolitos de las poblaciones de saboga, Alosa fallax (Lacépède, 1803), de los ríos Ulla y Miño (Doctoral thesis, Universidade de Santiago de Compostela, Spain). 10.13140/RG.2.2.18821.35044

Nachón, D. J., Bareille, G., Drouineau, H., Tabouret, H., Taverny, C., Boisneau, C., Berail, S., Pécheyran, C., Claverie, F., & Daverat, F. (2020). 1980s population-specific compositions of two related anadromous shad species during the oceanic phase determined by microchemistry of archived otoliths. Canadian Journal of Fisheries and Aquatic Sciences, 77(1), 164–176. 10.1139/cjfas-2018-0444

Nachón D. J., Mota M., Antunes C., Servia M. J., & Cobo F. (2016). Marine and continental distribution and dynamic of the early spawning migration of twaite shad (*Alosa fallax* (Lacépède, 1803)) and allis shad (*Alosa alosa* (Linnaeus, 1758)) in the north-west of the Iberian Peninsula. Marine and Freshwater Research, 67, 1229–1240. 10.1071/MF14243

Nachón, D. J., Pico, A., Vieira-Lanero, R., Barca, S., Cobo, M. d. C., & Cobo, F. (2022a). Analysis of Bycatches of Two Related Anadromous Shad Species in Fisheries along the Galician Atlantic Coast (NW Iberian Peninsula, Southwest Europe). Biology and Life Sciences Forum, 13(1), 57. 10.3390/blsf2022013057

Nachón, D. J., Pico, A., Vieira-Lanero, R., Barca, S., Cobo, M. d. C., & Cobo, F. (2022c). Biology and Ecology of Two Anadromous Species of the Genus *Alosa* (*A. alosa* and *A. fallax*) in the Galician Coastal Marine Environment Based on Bycatch Individuals: Proposals for the Improvement of Their Protection and Management. Biology and Life Sciences Forum, 13(1), 55. 10.3390/blsf2022013055

Nachón, D. J., Vieira-Lanero, R., Barca, S., Pico, A., Cobo, M. d. C., & Cobo, F. (2022b). Análisis de las capturas incidentales y de los descartes de dos especies anádromas del género *Alosa* (*A. alosa* y *A. fallax*) efectuados por la flota costera de Galicia. In M. Rey-Méndez, J. Fernández Casal, M. A. Lastres, N. González-Henríquez, & X. A. Padín (Eds.), Foro dos Recursos Mariños e da Acuicultura das Rías Galegas (pp. 173–179). Santiago de Compostela: Foroacui.

Nachón, D. J., Vieira-Lanero, R., Barca, S., Pico, A., Cobo, M. d. C., & Cobo, F. (2022d). Biología y ecología de dos especies anádromas del género *Alosa* (*A. alosa* y *A. fallax*) en el medio marino costero gallego a partir de los datos aportados por las capturas accidentales: propuestas para la mejora de su protección y gestión. In M. Rey-Méndez, J. Fernández Casal, M. A. Lastres, N. González-Henríquez, & X. A. Padín (Eds.), Foro dos Recursos Mariños e da Acuicultura das Rías Galegas (pp. 181–188). Santiago de Compostela: Foroacui.

Nachón, D. J., Vieira, R., & Cobo, F. (2019a). Sábalo – *Alosa alosa*. In P. López, J. Martín, & F. Cobo (Eds.), Enciclopedia Virtual de los Vertebrados Españoles. Museo Nacional de Ciencias Naturales, Madrid. Available at: http://www. https://www.vertebradosibericos.org/peces/aloalo.html (last accessed 15 April 2025).

Nachón, D. J., Vieira, R., & Cobo, F. (2019b). Saboga – *Alosa fallax*. In P. López, J. Martín, & F. Cobo (Eds.), Enciclopedia Virtual de los Vertebrados Españoles. Museo Nacional de Ciencias Naturales, Madrid. Available at: http://www. https://www.vertebradosibericos.org/peces/alofal.html (last accessed 15 April 2025).

Nakagawa, S., & Cuthill, I. C. (2007). Effect size, confidence interval and statistical significance: A practical guide for biologists. Biological Reviews, 82(4), 591–605. 10.1111/j.1469-185X.2007.00027.x

OSPAR (2009). Background Document for Allis Shad *Alosa alosa*. Biodiversity Series. Available at: http://qsr2010.ospar.org/media/assessments/Species/P00418_Allis_shad.pdf (last accessed 15 April 2025).

Poloczanska, E. S., Brown, C. J., Sydeman, W. J., Kiessling, W., Schoeman, D. S., Moore, P. J., Brander, K., Bruno, J. F., Buckley, L. B., Burrows, M. T., Duarte, C. M., Halpern, B. S., Holding, J., Kappel, C. V., ÓConnor, M. I., Pandolfi, J. M., Parmesan, C., Schwing, F., Thompson, S. A., & Richardson, A. J. (2013). Global imprint of climate change on marine life. Nature Climate Change, 3(10), 919–925. 10.1038/nclimate1958

Poulet, C., Barber-O’Malley, B. L., Lassalle, G., & Lambert, P. (2022). Quantification of land–sea nutrient fluxes supplied by allis shad across the species’ range. Canadian Journal of Fisheries and Aquatic Sciences, 79(3), 395–409. 10.1139/cjfas-2021-0012

Poulet, C., Lasalle, G., Jordaan, A., Limburg, K. E., Nack, C. C., Nye, J. A., ÓMalley, A., Barber-O’Malley, B. L., Stich, D. S., Waldman, J. R., Zydlewski, J., & Lambert, P. (2023). Effect of straying, reproductive strategies, and ocean distribution on the structure of American shad populations. Ecosphere, 14(2), e4712. 10.1002/ecs2.4712

Puncher, G. N., Cariani, A., Maes, G. E., Van Houdt, J., Herten, K., Cannas, R., Rodríguez-Ezpeleta, N., Albaina, A., Estonba, A., Lutcavage, M., Hanke, A., Rooker, J., Franks, J. S., Quattro, J. M., Basilone, G., Fraile, I., Laconcha, U., Goñi, N., Kimoto, A., Macías, D., Alemany, F., Deguara, S., Zgozi, S., Garibaldi, F., Oray, I., K., Karakulak, F., S., Abid, N., Santos, M. N., Addis, P., Arrizabalaga, H., & Tinti, F. (2018). Spatial dynamics and mixing of bluefin tuna in the Atlantic Ocean and Mediterranean Sea revealed using next-generation sequencing. Molecular Ecology Resources, 18(3), 620–638. 10.1111/1755-0998.12764

Randon, M., Daverat, F., Bareille, G., Jatteau, P., Martin, J., Pecheyran, C., & Drouineau, H. (2018). Quantifying exchanges of Allis shads between river catchments by combining otolith microchemistry and abundance indices in a Bayesian model. ICES Journal of Marine Science, 75(1), 9–21. 10.1093/icesjms/fsx148

Rougemont, Q., Perrier, C., Besnard, A. L., Lebel, I., Abdallah, Y., Feunteun, E., Réveillac, R., Lasne, E., Acou, A., Nachón, D. J., Cobo, F., Evanno, G., Baglinière, J. L., & Launey, S. (2022). Population genetics reveals divergent lineages and ongoing hybridization in a declining migratory fish species complex. Heredity, 129(2), 137–151. 10.1038/s41437-022-00547-9

Rougier, T., Lambert, P., Drouineau, H., Girardin, M., Castelnaud, G., Carry, L., Aprahamian, M. W., Rivot, E., & Rochard, E. (2012). Collapse of allis shad, *Alosa alosa*, in the Gironde system (southwest France): environmental change, fishing mortality, or Allee effect?. ICES Journal of Marine Science, 69(10), 1802–1811. 10.1093/icesjms/fss149

Sabatié, M. R. (1993). Recherches sur l’écologie et la biologie des aloses au Maroc (Alosa alosa Linné, 1758 et Alosa fallax Lacépède, 1803): exploitation et taxonomie des populations atlantiques, bioécologie des aloses de l’oued Sebou (Doctoral thesis, Université de Bretagne Occidentale, France).

Sabatino, S. J., Faria, R., & Alexandrino, P. B. (2022). Genetic structure, diversity, and connectivity in anadromous and freshwater *Alosa alosa* and *A. fallax*. Marine Biology, 169(1), 1–14. 10.1007/s00227-021-03970-4

Sánchez-Hernández, J., Gabler, H. M., & Amundsen, P. A. (2017). Prey diversity as a driver of resource partitioning between river-dwelling fish species. Ecology and Evolution, 7(7), 2058–2068. 10.1002/ece3.2793

Schlosser, I. J. (1998). Fish recruitment, dispersal, and trophic interactions in a heterogeneous lotic environment. Oecologia, 113(2), 260–268. 10.1007/s004420050377

Smoliński, S., Schade, F. M., & Berg, F. (2020). Assessing the performance of statistical classifiers to discriminate fish stocks using Fourier analysis of otolith shape. Canadian Journal of Fisheries and Aquatic Sciences, 77(4), 674–683. 10.1139/cjfas-2019-0251

Stratoudakis, Y., Antunes, C., Correia, C., Belo, A. F., & Almeida, P. R. (2023). Riverine communities and management systems for anadromous fisheries in the Iberian Peninsula: global strategy, local realities. Reviews in Fish Biology and Fisheries, 33(3), 875–892. 10.1007/s11160-022-09742-7

Stratoudakis, Y., Correia, C., Belo, A. F., & Almeida, P. R. (2020). Improving participated management under poor fishers’ organization: Anadromous fishing in the estuary of Mondego River, Portugal. Marine Policy, 119, 104049. 10.1016/j.marpol.2020.104049

Stratoudakis, Y., Mateus, C. S., Quintella, B. R., Antunes, C., & Almeida, P. R. (2016). Exploited anadromous fish in Portugal: suggested direction for conservation and management. Marine Policy, 73, 92–99. 10.1016/j.marpol.2016.07.031

Taillebois, L., Sabatino, S., Manicki, A., Daverat, F., Nachón, D. J., & Lepais, O. (2020). Variable outcomes of hybridization between declining *Alosa alosa* and *Alosa fallax*. Evolutionary applications, 13(4), 636–651. 10.1111/eva.12889

Taverny, C. (1991). Contribution à la connaissance de la dynamique des populations d’aloses (Alosa alosa L. et Alosa fallax Lacépède), dans le système fluvio-estuarien de la Gironde: pêche, biologie et écologie. Étude particulière de la dévalaison et de l’impact des activités humaines (Doctoral thesis, Université de Bordeaux 1, France).

Taverny, C., Belaud, A., Elie, P., & Sabatié, M. R. (2000a). Influence des activités humaines. In J. L. Baglinière & P. Elie (Eds.), Les aloses (Alosa alosa et Alosa fallax spp.). Écobiologie et variabilité des populations (pp. 227–248). Paris: INRA-CEMAGREF.

Taverny, C., Cassou-Leins, J. J., Cassou-Leins, F., & Elie, P. (2000b). De l’œuf à l’adulte en mer. In J. L. Baglinière & P. Elie (Eds.), Les aloses (Alosa alosa et Alosa fallax spp.). Écobiologie et variabilité des populations (pp. 93–124). Paris: INRA-CEMAGREF.

Taverny, C., & Elie, P. (2001). Répartition spatio-temporelle de la grande alose *Alosa alosa* (Linné, 1766) et de ĺalose feinte *Alosa fallax* (Lacépède, 1803) dans le Golfe de Gascogne. Bulletin Français de la Pêche et de la Pisciculture, 362/363, 803–821. 10.1051/kmae:2001020

Thorrold, S. R., Latkoczy, C., Swart, P. K., & Jones, C. M. (2001). Natal homing in a marine fish metapopulation. Science, 291(5502), 297–299. 10.1126/science.291.5502.297

Trancart, T., Rochette, S., Acou, A., Lasne, E., & Feunteun, E. (2014). Modeling marine shad distribution using data from French bycatch fishery surveys. Marine Ecology Progress Series, 511, 181–192. 10.3354/meps10907

Verhelst, P., Reubens, J., Buysse, D., Goethals, P., Van Wichelen, J., & Moens, T. (2021). Toward a roadmap for diadromous fish conservation: the Big Five considerations. Frontiers in Ecology and the Environment, 19(7), 396–403. 10.1002/fee.2361

Walther, B. D., & Thorrold, S. R. (2006). Water, not food, contributes the majority of strontium and barium deposited in the otoliths of a marine fish. Marine Ecology Progress Series, 311, 125–130. 10.3354/meps311125

Walther, B. D., & Thorrold, S. R. (2008). Continental-scale variation in otolith geochemistry of juvenile American shad (*Alosa sapidissima*). Canadian Journal of Fisheries and Aquatic Sciences, 65(12), 2623–2635. 10.1139/F08-164

Walther, B. D., & Thorrold, S. R. (2010). Limited diversity in natal origins of immature anadromous fish during ocean residency. Canadian Journal of Fisheries and Aquatic Sciences, 67(10), 1699–1707. 10.1139/F10-08

Walther, B. D., Thorrold, S. R., & Olney, J. E. (2008). Geochemical signatures in otoliths record natal origins of American shad. Transactions of the American Fisheries Society, 137(1), 57–69. 10.1577/T07-029.1

Wilson, K., & Veneranta, L. (2019). Data-limited diadromous species – review of European status. ICES Cooperative Research Reports (CRR) No. 348. Available at: https://ices-library.figshare.com/articles/report/Data-limited_diadromous_species_review_of_European_status/18624041?file=33403097 (last accessed 16 April 2025).

Zimmerman, C. E., Swanson, H.K., Volk, E.C., & Kent, A. J. (2013). Species and life history affect the utility of otolith chemical composition for determining natal stream of origin for Pacific salmon. Transactions of the American Fisheries Society, 142(5), 1370–1380. 10.1080/00028487.2013.811102

